# Open Force Field Consortium: Escaping atom types using direct chemical perception with SMIRNOFF v0.1

**DOI:** 10.1101/286542

**Authors:** David L. Mobley, Caitlin C. Bannan, Andrea Rizzi, Christopher I. Bayly, John D. Chodera, Victoria T. Lim, Nathan M. Lim, Kyle A. Beauchamp, Michael R. Shirts, Michael K. Gilson, Peter K. Eastman

**Author notes:** **For correspondence:** (DLM).

## Abstract

Here, we focus on testing and improving force fields for molecular modeling, which see widespread use in diverse areas of computational chemistry and biomolecular simulation. A key issue affecting the accuracy and transferrability of these force fields is the use of atom typing. Traditional approaches to defining molecular mechanics force fields must encode, within a discrete set of atom types, *all* information which will ever be needed about the chemical environment; parameters are then assigned by looking up combinations of these atom types in tables. This atom typing approach leads to a wide variety of problems such as inextensible atom-typing machinery, enormous difficulty in expanding parameters encoded by atom types, and unnecessarily proliferation of encoded parameters. Here, we describe a new approach to assigning parameters for molecular mechanics force fields based on the industry standard SMARTS chemical perception language (with extensions to identify specific atoms available in SMIRKS). In this approach, each force field term (bonds, angles, and torsions, and nonbonded interactions) features separate definitions assigned in a hierarchical manner without using atom types. We accomplish this using *direct chemical perception*, where parameters are assigned directly based on substructure queries operating on the molecule(s) being parameterized, thereby avoiding the intermediate step of assigning atom types — a step which can be considered *indirect* chemical perception. Direct chemical perception allows for substantial simplification of force fields, as well as additional generality in the substructure queries. This approach is applicable to a wide variety of (bio)molecular systems, and can greatly reduce the number of parameters needed to create a complete force field. Further flexibility can also be gained by allowing force field terms to be interpolated based on the assignment of fractional bond orders via the same procedure used to assign partial charges. As an example of the utility of this approach, we provide a minimalist small molecule force field derived from Merck’s parm@Frosst (an Amber parm99 descendant), in which a parameter definition file only «300 lines long can parameterize a large and diverse spectrum of pharmaceutically relevant small molecule chemical space. We benchmark this minimalist force field on the FreeSolv small molecule hydration free energy set and calculations of densities and dielectric constants from the ThermoML Archive, demonstrating that it achieves comparable accuracy to the Generalized Amber Force Field (GAFF) that consists of many thousands of parameters.

## 1 Introduction

Classical all-atom molecular mechanics force fields see widespread use in molecular simulations in diverse areas in chemistry, biochemistry, biology, drug discovery, and materials science [1-9]. Often, these are two-body additive fixed-charge force fields with the relatively simple Lennard-Jones functional form for nonpolar interactions. Despite this simplicity, force fields have achieved remarkable successes calculating and even predicting a wide range of properties and behaviors far beyond the simple gas phase [1], condensed phase [1, 10] or biomolecular [9, 11-13] properties for which they have been parameterized. Essentially, force fields take a complex parameter optimization problem and reduce the complexity considerably by relying on reasonable approximations of the underlying physics in order to reduce the amount of data required to produce force fields with broad applicability while also simplifying the parameter fitting process. Thus, force fields demonstrate their considerable success even in blind predictions, such as on host-guest [14-16] and protein-ligand binding affinities [17-26], hydration free energies [27, 28], partition and distribution coefficients [29], ligand binding modes and activity [30], and others. Extensive retrospective tests for other properties such as dielectric constants [31-34] and perturbations in protein stability [35] are also worth noting given their success. It seems safe to say these relatively simple models have succeeded far beyond original expectations, likely in part because of their strong physical basis, careful and time-consuming attention paid in parameterization, and a reasonable balance of speed versus accuracy for many applications.

At the same time, force fields also have numerous clear failure cases indicating further need for improvement. For example, in the present day GAFF [36] and GAFF2 force fields, certain aromatic rings can buckle due to human error resulting from complexities of atom typing (Figure 1).

**Figure 1.**
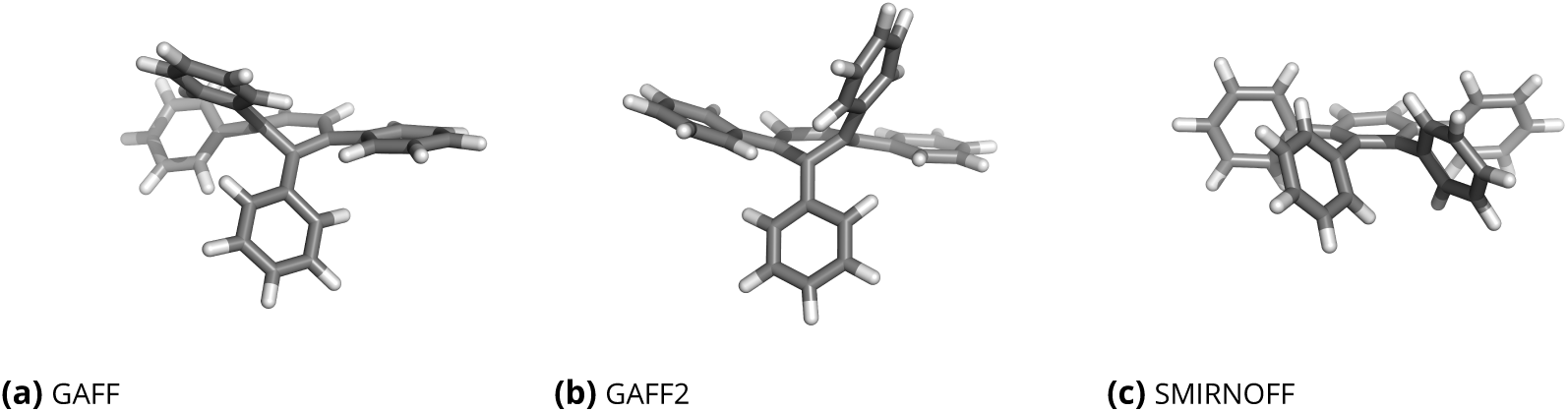
Representative geometries of 1,2,3,4-tetraphenylbenzene. Depending on what force field is applied, the typical geometry of 1,2,3,4-tetraphenylbenzene can have an incorrect buckling of the central aromatic ring in molecular dynamics simulations. Here, we show representative geometries from short gas-phase simulations with each force field. In GAFF and GAFF2 (a-b), the central ring buckles, whereas with the prototype force field we introduce here, SMIRNOFF99Frosst, the aromatic ring remains planar as expected. SMIRNOFF99Frosst recognizes the aromatic nature of these torsions using a torsional parameter applied using the SMIRKS pattern `[*:1]~ [#6X3:2]:[#6X3:3] ~ [*:4]` (see Table 1), whereas the `parmchk2` tool used to infer the torsional parameters needed for use with GAFF/GAFF2 incorrectly assigns some of the aromatic bonds within the central ring torsions which are appropriate for rotatable single bonds, ultimately resulting in buckling. As noted in Figure 2d, atom typing in this case is complex.

One key concern in developing force fields is the balance of accuracy versus generality. Since the underlying functional form is certainly approximate, it is always possible to improve accuracy by adding more parameters which are tuned to particular use cases. Of course, in some cases these additional parameters are warranted; for example, a tetrahedral carbon obviously requires a different bonding geometry than a planar one and any model missing this will result in major structural errors. However, in other cases specialization may be unwarranted or even lead to problems of transferability due to overfitting. Typically, the major, general-purpose force field families used for molecular simulations have tried to achieve reasonable balance in this regard, in many respects hitting a sweet spot which provides sufficient chemical sophistication to ensure adequate coverage of and accuracy for major distinct chemical functionalities, while also not adding additional unnecessary parameters that might result in overfitting and perhaps impair generality.

### 1.1 Force field development usually requires a tremendous investment of human expertise and time

One major challenge in force field development is the amount of human time and expertise involved. Development of a new general force field (covering all or almost all of normal organic chemistry and biomolecules) from scratch typically takes many years, based on historical precedent. Thus, while there have been numerous adjustments to biomolecular force fields over the years, especially terms relating to proteins and nucleic acids [9, 11-13, 37], the core of most of our present-day force fields, at least aside from the torsions and charges, still seems to typically date to the 1980s and early 1990s. Building a new fixed-charge force field from scratch would simply require too much effort over too long a time for individual academic groups to tackle the problem, not to mention the fact that funding is difficult to impossible to obtain for such an academic effort. Small molecule force fields have thus received much less attention than biomolecular force fields, and have typically been developed at least in part by generalizing biomolecular force fields to cover more chemical space [36, 38, 39]. Thus development of small molecule force fields has lagged behind that of protein force fields, partly because of the additional chemical complexity involved and corresponding additional human time required.

### 1.2 We have not yet reached the accuracy limit for fixed-charge force fields

At the same time, fixed charge force fields show clear room for improvement. Certainly not all accuracy problems are due to force field problems, but it seems increasingly clear that results of calculations often are quite sensitive to force field parameters [40-44], and that force field issues do result in significant systematic errors in a fairly wide range of cases (e.g. [19, 42, 44, 45]). Indeed, in some cases, systematic errors can be traced back to problems with force field parameters for particular functional groups [46-48] and targeted follow-up work can in some cases fix these issues. For example, GAFF parameters yielded systematic errors for alkenes which could be fixed by a minor adjustment to Lennard-Jones parameters [46], and larger errors for alcohols in general due to issues with underpolarization of the hydroxyl group which were fixed by a focused effort [31]. Issues with certain bridgehead atoms persist to this day and could also likely be fixed via similar efforts (e.g. Figures 1 and 2e). But these isolated efforts serve as band-aids rather than a systematic fix, and are themselves human-intensive.

The evidence seems clear that a new generation of fixed-charge force fields developed from scratch could do dramatically better than our current force fields, but the investment of human time and expertise required is so large that no general effort has gone forward to refit force fields from scratch. OPLS3 [9] probably represents the largest recent fixed-charge force field effort, but even this relies on nonbonded parameters originally developed much earlier, in the 1980s [49] and 1990s [10]. COMPASS II also results from a large development effort but this, too, marks an extension of earlier work rather than an overall refitting [8]. In our view, the solution to this problem is to allow force fields to be fit with minimal human intervention. This could be achieved by automating the force field development process so that human expertise is used to select only the functional form and input data for parameterization, but then a completely automated machinery produces the force field itself. The process of replacing human-annotated featurizations with fully automated ones has recently led to advances in both chemistry (e.g., DeepChem [50]) and image recognition (e.g., ImageNet [51]), though we are not yet aware of applications to force field development.

### 1.3 Human expertise is used to decide atom types which are used to assign parameters via indirect chemical perception

While a reasonable amount of effort has gone into improving the automated fitting of parameters for a particular force field given input data, such as via ForceBalance [52-56], this approach still requires a great deal of human expertise deciding which parameters need to be fitted when building general purpose force fields. Notably, a human expert must decide how many atom types (and thus how many bond, angle, torsional, Lennard-Jones and charge parameters) are needed to represent all of the relevant chemistry, and then, given these choices and others, automated machinery can improve the numerical parameters associated with these force field terms. Atom typing rapidly becomes extremely complex even for relatively simple molecules, as illustrated in Figure 2. Thus this process becomes so arduous that some approaches instead attempt to build unique parameter sets for each individual molecule from scratch [57-60]. While this may be an interesting approach in some cases, we remain very interested in general purpose force fields that can be rapidly applied to large libraries of molecules, for which automation of force field development is critical.

**Figure 2.**
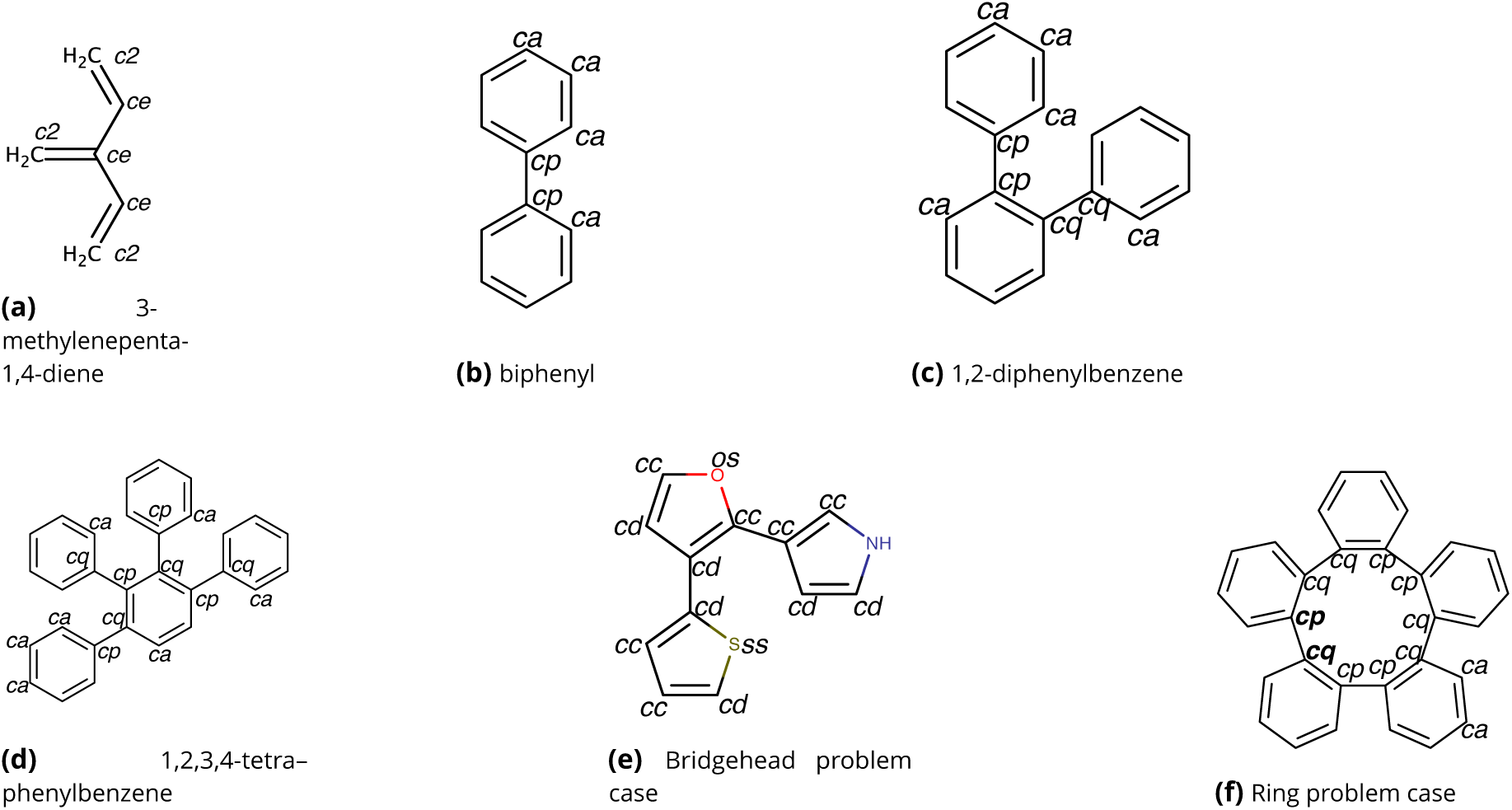
Even simple molecules can present challenging cases for indirect chemical perception via atom typing. Atom typing these molecules in a way which can lead to proper assignment of parameters can be challenging because of bonding patterns. Indicated atom types are GAFF/GAFF2 types. **In (a)**, all carbons are sp^2^ yet are connected by alternating single and double bonds, forcing introduction of the `ce` atom type for the inner sp^2^ carbons to allow single and double carbon-carbon bonds to be distinguished. **In (b)**, the bridgehead aromatic carbons in biphenyl must have a single bond joining them, forcing introduction of the `cp` atom type which is identical to the normal aromatic carbon (`ca`) except that `cp-cp` bonds are single and thus rotatable. Introduction of an additional phenyl ring, in 1,2-diphenylbenzene **(c)** further complicates matters since, with only the `ca` and `cp` types there would be a `cp-cp` (single) bond within the lower aromatic ring. GAFF/GAFF2 address this by introduction of a new type, `cq`, which is identical to `cp` and `ca` so that `cp-cp` bonds are single, `cq-cq` bonds are single, but `cq-cp` bonds (and bonds involving `ca` of any sort) are aromatic. This scheme can in principle handle even larger molecules like 1,2,3,4-tetraphenylbenzene **(d)** though, as discussed in the text, the massive proliferation of torsional parameters resulting from the numerous atom types employed leads to considerable potential for human error. However, this approach of introducing additional duplicate atom types requires a human expert to notice when such types will be needed. **(e)** shows a case where bridgehead bonds between five membered rings are typed as `cd-cd` or `cc-cc`, the same as bonds within aromatic rings. Thus, as we discussed in Section 4.4, GAFF/GAFF2 make these bonds non-rotatable, even though they are single bonds. This could presumably be fixed in the same manner as the issues of (b) and (c) via introduction of additional atom types. However, some molecules, like **(f)**, are impossible to atom type properly in this framework [36]; incorrect atom types are shown in boldface and will lead to misassignment of parameters.

To automate the entire force field generation process, we need to reduce the human expertise required even in early stages, such as atom typing. Specifically, our goal in this work is to eliminate predefined atom types and instead move to a chemical perception language which can be adjusted as *part* of a force field development process, paving the way for further work which will automate force field generation.

Atom typing can be thought of as a type of *indirect chemical perception* (Figure 3), where a molecule or molecules are processed via some machinery to assign labels to atoms (atom types) and then these labels are subsequently processed to assign parameters. Thus, key for success is ensuring that the atom types encode all of the relevant information but no *unnecessary* information, as once parameterization is begun, the atom typing rules are considered fixed. Subsequent addition of new atom types—for example, to extend the force field into new areas chemical space—creates enormous difficulties in how existing parameters should be adjusted to accommodate the need to fit newly created parameters [8] (though hierarchical schemes can assist with this, e.g. [62], and thus a hierarchical scheme provides part of our solution as well). Consequently, errors are frequent; to give just one example, GAFF/GAFF2 recognize certain single bonds between bridgehead atoms as non-rotatable due to oversights in designing atom typing (Figure 2e, Section 2.2).

**Figure 3.**
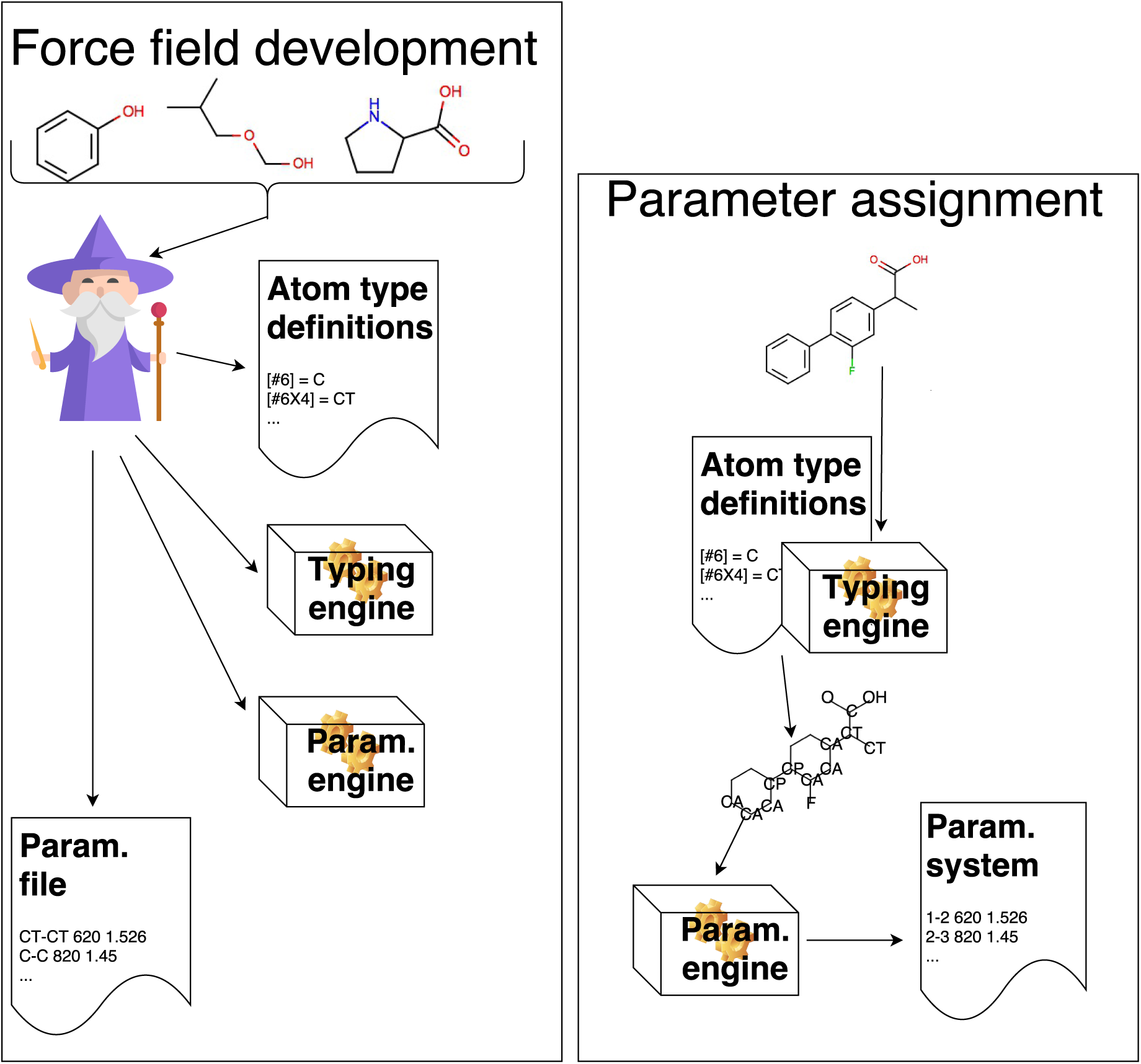
Indirect chemical perception requires that a large library of atom types encode all potentially relevant chemical environments. Force field assignment via indirect chemical perception requires several stages of processing. First, in the *force field development process (left)* a human expert (“wizard”) considers a set of molecules which the force field should cover and decides which chemical environments will be important to treat separately, choosing a set of atom types to bin this chemistry and tabulating or encoding these atom type definitions. The expert then encodes a typing engine which can assign these atom types to arbitrary molecules, writing out a chemical graph with atoms (nodes) labeled by atom types. Once this engine is in place, the expert separately encodes a parameterization machinery which will read in labeled chemical graphs and assign force field parameters based on atom types [61], often from a lookup table called a parameter file. This engine will write out the result to a file containing a parameterized system suitable for simulations. The expert also develops the parameter file which will be used by the parameterization engine. Second, in the *parameter assignment process* (right), a specific molecule or system is input into the typing engine previously developed, which applies the atom type definitions and writes out a labeled chemical graph. This labeled graph is then processed by the parameterization engine to produce a parameterized system suitable for simulation. This process is indirect—the parameterization engine considers a labeled graph, not the molecule itself. Thus, in this final step, all of the relevant information about distinct chemical environments must be encoded by the atom types and other information in the graph (in AMBER-family force fields, just the atom types and connectivity). (Wizard icon made by Freepikfrom www.flaticon.com)

Force field parameters could instead be assigned by a process of *direct chemical perception*, which might bypass atom typing altogether and would assign parameters to individual atoms, bonds, angles, or torsions by processing the full molecular graph (consisting of all atoms in their standard valence representation, with connectivity, bond orders, and an aromaticity model applied) directly via a chemically-aware engine. To see the distinction, note that force fields in the AMBER force field family do not retain bond order when assigning parameters, so if any bond order information is necessary, this must be encoded in the labels or atom types themselves, as we discuss further below. In contrast, a tool doing direct chemical perception could use information about a molecule such as bond order, as it operates directly on the molecular graph itself rather than an intermediate labeled graph that no longer retains bond orders or access to other molecular properties.

In some cases the distinction between indirect and direct chemical perception can become somewhat blurry, as we discuss further below (Section 2.2.1), especially when force fields employing indirect chemical perception begin making extremely sophisticated atom types to attempt to deal with anticipated parameterization problems.

## 2 How we got to where we are

### 2.1 Why force fields often use indirect chemical perception

Current biomolecular simulation packages largely use indirect chemical perception, but deciphering exactly *why* from the literature is nontrivial. As best we can tell, two major factors led to the current state of affairs - one a matter of early research focus, and another a matter of software architecture.

First, the early development of force fields focused largely on specific molecules or use cases where the *use* of parameters posed a more important, interesting, and pressing scientific problem than how to *assign* parameters in general. Thus much early work focused on either a limited number of small organic molecules where parameter assignment could easily be done by hand, or on biopolymers which are composed of a limited number of basic building blocks of well-defined chemistry [63-70]. In either case the problem of chemical perception is not a major factor.

Second, the first general force fields for organic molecules which are still in widespread current use came as late as the mid-1990s (especially GAFF [36] and MMFF94 [71-73]) by which time a number of simulation packages were already well established and fairly tied to indirect chemical perception, where atom types are first assigned and then used to look up parameters. This meant that when Antechamber was developed to automatically assign atom types for AMBER’s GAFF small molecule force field (probably making AMBER the first academic biomolecular force field to have an automatic tool for general small molecule paramerization [74]), its developer could state:

> “It is often believed that bond types (single, double, triple, aromatic single, aromatic double, conjugated single and conjugated double) are essential to precisely describe a chemical environment. However, for some reasons most of biological molecule force fields, including AMBER, only apply atom types in the force field parameter files. The bond type information is also sacrificed in GAFF for consistency with the AMBER force fields and the programs of the AMBER packages.” [38]

The authors go on to note that, because of the lack of bond type information available to the software, a variety of new atom types had to be introduced to attempt to capture this chemistry, and they also highlight cases where atom types simply *cannot* capture all of the requisite details. Discussions with one of the original authors highlighted that doing otherwise would have required too large a change to the underlying software and was thus deemed impractical [75]. While MMFF94 seems to have avoided some of the problems encountered by GAFF, it, too, uses indirect chemical perception [71, 72]. Many simulation packages seem to have been hampered by a similar process to that encountered by AMBER’s GAFF and Antechamber — they selected file formats and data structures which do not retain bond order information or other aspects of the underlying chemistry needed for direct chemical perception, essentially *requiring* force fields to rely on atom types. This leads to pathologies handling conjugated bonds that have been propagated to contemporary force fields, exacerbating other difficulties of atom typing.

Indirect chemical perception is so common that it is almost deemed essential to force fields; for example, the developers of CGenFF, the CHARMM general force field, remark, “Thus, the first step of assigning parameters for a chemical system is assigning atom types to that system.” [74] Thus, while the chemical perception used by CGenFF does attempt to go past that employed by Antechamber, it still uses indirect chemical perception, first assigning atom types and then using these to assign parameters (thus it, too, runs into instances of molecules which cannot be typed). OPLS3 also maintains a reliance on indirect chemical perception [9].

Thus, it appears that the field did not make a deliberate decision to employ indirect chemical perception because it was superior, but instead has gone that route for historical and architectural reasons. At the same time, the literature has also highlighted deficiencies of this approach, such as un-typeable molecules for GAFF and CGenFF [38, 74], and the fact that force fields employing indirect chemical perception simply *cannot* recognize any differences in chemistry that are not encoded by atom types [76-78].

This discussion is not to imply that indirect chemical perception is necessarily inferior; indeed, some work with indirect chemical perception uses quite sophisticated chemical perception [38, 72, 74, 78]; rather, the larger point is that direct chemical perception allows greater flexibility for force field developers.

### 2.2 Atom typing introduces a range of problems

While, in our view, atom typing has been extremely helpful in developing general and relatively transferable force fields which have allowed a great deal of progress in applications of molecular modeling (MMFF94 [71, 72], GAFF [36], and CGenFF [74] have been crucial in enabling widespread modeling of protein-ligand interactions), we believe it has also impaired force field science and force field development and we hope to change that. Atom typing requires a human expert, and poses an arduous task [8] introducing “a certain degree of ambiguity and arbitrariness” [79]. Given the expertise required, then, most work on general purpose (bio)molecular force fields is done by select individuals in just a handful of groups. The chemical perception for atom typing is typically hard coded into software tools where it is often both invisible and hard to modify (though there are efforts to change this [80]). Overall, this impairs force field science because very few individuals or groups have the necessary expertise to modify, extend, or even troubleshoot the available expert systems for atom typing, though some efforts are being made to improve this, such as by hierarchical atom typing [62].

Atom typing also provides a key place where early decisions or even mistakes can lead to subsequent problems for force field development which are hard to overcome. For one, atom typing makes an up-front decision as to how to bin chemical space. Once separate atom types are assigned, it is difficult to bin chemical space differently, even if the data might warrant it. Additionally, an introduction of a new atom type to fix a problem with one valence term results in a proliferation of parameters for all valence terms. To apply automated parameterization machinery when many equal parameters exist (such as the 16 sets of Lennard-Jones parameters for carbon in GAFF/GAFF2 which only have three distinct values [39]), a human expert would have to designate which parameters should be constrained to be identical versus which should be allowed to be distinct.

#### 2.2.1 Specific problem molecules help illustrate the challenges faced when atom typing

For example, in 3-methylenepenta-1,4-diene (Figure 2a), atom typing is difficult to implement in a general way. The carbon atoms involved all have similar chemical environments, with one double bond to another atom and single bonds to three other atoms. The differences are simply in how many hydrogen atoms they are bonded to. Yet the resulting atom typing must have sufficient complexity to encode the fact that bond orders between carbons are alternating single and double. AMBER family force fields deal with this by introducing additional atom types to essentially encode bond order information (GAFF atom types are shown in the figure). In GAFF, Lennard-Jones parameters for the different types used are identical and bond parameters are virtually identical for the carbon-hydrogen single bonds involved (force constants 342.5 kcal/(mol Å^2^) with length 1.0883 Å for `c2-ha`, 344.3 kcal/(mol Å^2^ and length 1.0870 Å for `ce-ha`). Direct chemical perception bypasses this complexity by simply applying the same nonbonded and bonded parameters to all of the carbons in this molecule except where differences are warranted (such as for rotation around single bonds versus double bonds and for geometries of some of the angles involved).

The biphenyl, 1,2-diphenylbenzene, and related series of molecules also provides an interesting challenge (Figure 2b-2d). In the simplest approach to atom typing, all the aromatic carbons would get a single atom type (ca in GAFF). However, as a consequence of indirect chemical perception, it is difficult to recognize the single bond between phenyl rings from a straightforward application of an “aromatic carbon” atom type to all of the ring atoms, so a new atom type for bridgehead carbon atoms was introduced [36, 81], `cp` in GAFF. This fixes the immediate issue, but in 1,2-diphenylbenzene (Figure 2c) there is an aromatic bond between two bridgehead carbon atoms. This can be again fixed by introducing a new atom type, `cq` in GAFF, which is identical to `cp` except that `cp-cq` bonds are single. This results in a proliferation of atom types (and requisite bond, angle, and torsional parameters) simply to work around a problem which would trivially be solved by using bond orders when assigning parameters. It also obfuscates the underlying chemistry—the relevant GAFF atom types (`ca`, `cp`, and `cq`) are only different because of the bond order of the atoms involved, but this would not be obvious to a non-expert practitioner. We will revisit this case later (4.4) to see how direct chemical perception deals differently with such cases.

While the biphenyl case has received careful attention to avoid incorrectly treating the bond between bridgehead carbons, this approach requires a human expert to identify likely problem cases and thus fails when encountering chemistry which has not received such careful attention. For example, in testing, we found that a similar scenario occurs for bonds between the GAFF/GAFF2 types `cc-cc`, `cc-cd`, and `cd-cd`, which are identical and are used for carbons in non-pure aromatic systems (Figure 2e). GAFF/GAFF2 give the torsions for these single bonds a barrier height of 16.00 kcal/mol, identical to the aromatic bonds within the five membered rings and even higher than the 14.5 kcal/mol barrier height used for aromatic bonds in biphenyl. Thus unexpected bridgehead atoms result in a rotatable single bond being treated as aromatic.

Such issues with complex atom typing also result in molecules which are impossible to atom type correctly [36] (Figure 2f) using the present AMBER machinery (which does not have access to bond order information), and which may also pose challenges for other force fields.

To help understand these issues, it is worth noting that indirect chemical perception, as we define it here, assigns atom types purely on the basis of their chemical environment in the molecule (where this environment can be as large as needed but is typically relatively local) and then other parameters are assigned based on the connectivity graph with vertices labeled using atom types, as illustrated in Figure 3. Figure 2 illustrates some cases where this approach fails because, for example, in Figure 2d, all atoms labeled `cp` and `cq` are in identical environments and therefore must be assigned identical types. GAFF fixes this by introducing new atom types for atoms with equivalent environments, in anticipation of the fact that these will need to be assigned different parameters. In some sense this procedure is already a step towards direct chemical perception, but inserted into a parameter assignment machinery expecting indirect chemical perception; rather than directly assigning bond types, GAFF attempts to encode the bond types into the atom types and use the existing machinery to assign parameters.

#### 2.2.2 Atom typing introduces considerable potential for human error

Despite the complexities of atom typing, humans do attempt to extend force fields by hand, and there, atom typing poses further challenges and introduces considerable potential for human error. For example, AMBER-family force fields introduced the H1, H2, and H3 atom types because of the apparent need for additional Lennard-Jones parameters for hydrogen [82]. However, introducing these parameters also requires a significant number of new torsions and, as we find below, the opportunity for human error to creep in where these torsions are simply omitted. The reverse also occurs, in that introducing new torsion types (as in the biphenyl family cases of Figure 2) results in extra Lennard-Jones parameters. Thus, a simple “fix” to one type of parameter in the force field can easily lead to human error elsewhere, such as omitted or erroneous torsional parameters.

The complexity of atom typing (and the lack of independence of the chemical perception for different parameter types) also makes parameterization vastly more complicated. For example, AMBER16’s GAFF 1.8 has 6,387 lines of parameters and GAFF2 (version 2.1) has 6,796. A key question for applying automated methods like ForceBalance is how many of these constitute parameters which should be fitted separately, versus how many should in fact be equivalent (like Lennard-Jones parameters for carbons in biphenyl) but are different only because of indirect chemical perception. One concrete example of this issue is that the `CA-CA` carbon aromatic bond in AMBER’s parm96 [69, 83] and parm99 sets [81] has a length of 1.40 Å whereas `CA-CB` (between substantially the same types except that `CB` is an an aromatic carbon at the junction of 5- and 6-membered rings, such as in adenosine and tryptophan) has a length of 1.404 Å, differing by only 0.004 Å, with no clear data indicating that this difference is warranted or truly significant [83] given the precision of the calculations. (Force constants for these bonds are identical.)

### 2.3 We propose a shift to direct chemical perception

Here, we propose a shift away from atom typing to direct chemical perception (DCP) and explain how it can help resolve many of the above problems with atom typing, as well as open up new lines of inquiry. We introduce direct chemical perception in general, explain how it can help simplify force fields and make them more extensible, introduce a new force field format using direct chemical perception, and provide a prototype new small molecule force field which uses direct chemical perception.

## 3 Theory of direct chemical perception

### 3.1 Direct chemical perception (DCP) assigns force field parameters directly to parts of molecules

DCP allows direct assignment of parameters via processing the full chemical graph of molecules, avoiding the limitations of atom types (Figure 4) — instead of assigning parameters by processing a connectivity graph labeled with predefined atom types, DCP can assign parameters via operations acting directly on the full chemical or molecular graph. For our purposes purpose, the “full chemical graph” here is defined as the standard valence bonded representation of the molecule with explicit hydrogens, formal charges, and an aromaticity model applied. Essentially, DCP involves means using a chemical perception language to assign force field parameters based on molecular fragments. DCP can be used to encode traditional atom type-based force fields by encoding the same chemical perception, so it can even reproduce pathologies or complexities associated with atom typing (with enough effort) such as the complex treatment of bridgehead atoms in GAFF’s handling of biphenyls. Direct chemical perception avoids these problems naturally. For example, the bond between aromatic rings in biphenyl is a single bond, and thus DCP can easily recognize it as requiring different bonded parameters than the aromatic bonds within the aromatic rings. Additionally, since the parameterization engine has access to the molecular graph, it has bond order information as well as full access to all information about the chemical environment. Thus, a variety of tools can be applied in parameterization, including (if needed) electronic structure calculations.

**Figure 4.**
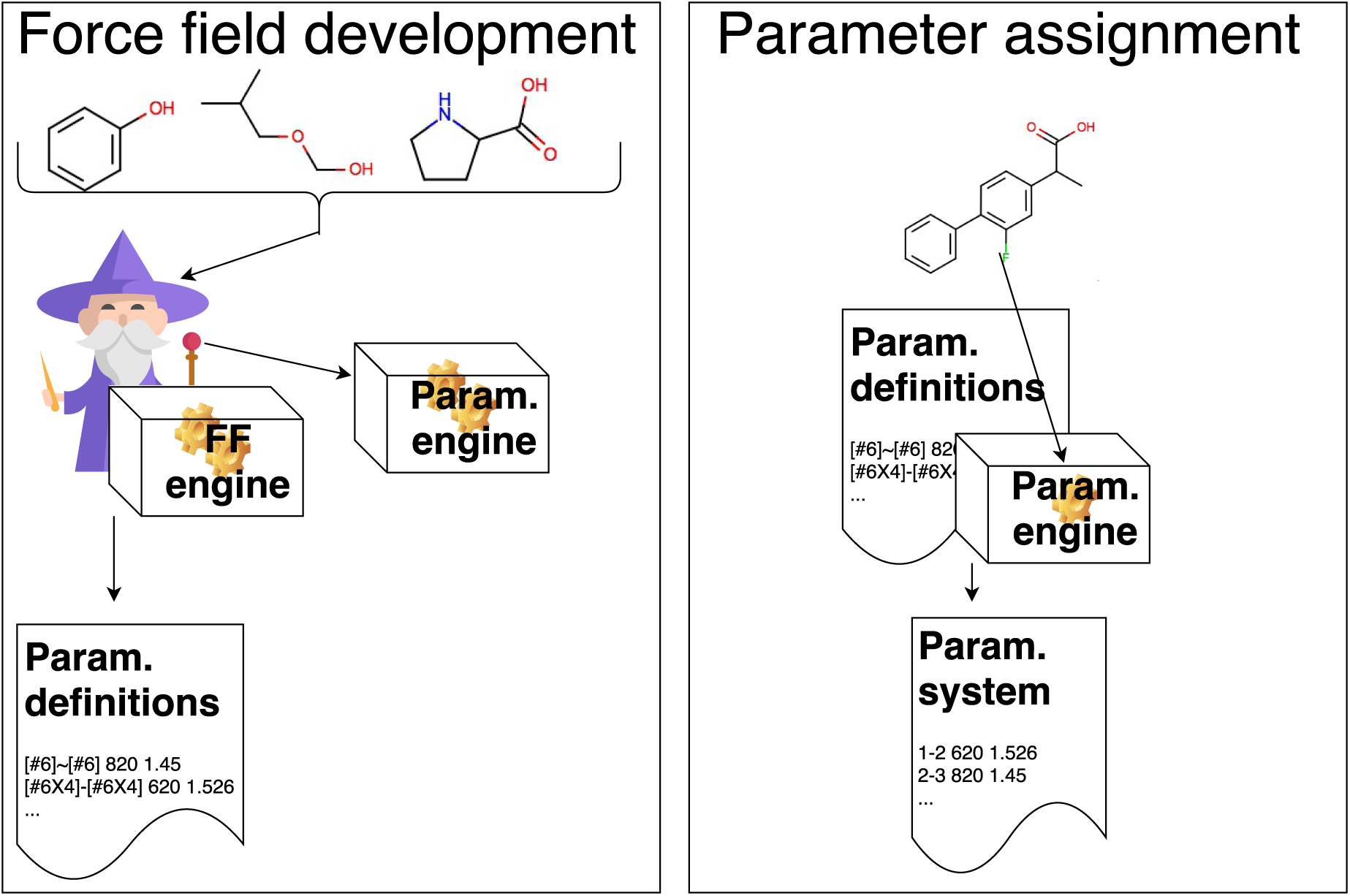
Direct chemical perception eliminates the need to encode all relevant chemical environment information in arbitrary predefined atom types. Force field assignment via direct chemical perception works on the full chemical graph of the molecules involved (including elements, connectivity, bond order, etc.), rather than first encoding information about the chemical environment into a complex set of predetermined atom types. First, in the *force field development process* (left) a human expert (“wizard”) and/or an automated method (a force field engine, “FF engine”) considers a set of molecules which the force field should cover (as well as potentially input data) and develops a force field to cover this chemistry, producing a set of parameter definitions and a parameterization engine that can apply these to molecules. Second, in the *parameter assignment process* (right), a specific molecule or system is input into the parameterization engine previously developed, which processes the molecule and uses the parameter definitions to apply force field parameters, producing a parameterized system suitable for simulation. The parameter assignment process is direct—the parameterization engine acts directly on the chemical graph of the molecules comprising the system, so all chemical environment information provided (or computable) is available to the engine. Unlike indirect chemical perception, there is no intermediate step of assigning atom type labels to a molecular graph; parameters are assigned directly based on the chemistry. (Wizard icon made by Freepik from www.fiaticon.com)

### 3.2 DCP avoids obfuscating chemistry and provides greater extensibility

Direct chemical perception avoids hiding the chemistry addressed by individual terms of the force field under an additional layer of encoding which can obscure the intent. For example, consider parameter assignment for valence parameters within ring systems of various sizes. In today’s fixed charge force fields, bond stretch parameters within such rings are dominated by the order (single, aromatic, or double) of the bonds involved, with modulation in some cases by the number of attached electron withdrawing or donating groups, but the *size* of the rings involved plays very little role in the bond stretch parameters. In contrast, angle bending parameters for the same rings show almost the exact opposite behavior - bond order matters comparatively little because the angle is primarily dictated by the geometry of the ring, whereas the size of the ring plays a huge role in determining the equilibrium angle. Thus, it seems that one type of chemical perception, focused primarily on bond order, is appropriate for assigning bond stretching parameters, whereas another is more appropriate for angle bending parameters.

However, indirect chemical perception typically applies *the same chemical perception for all force types*, so if we need to introduce new atom types to capture the correct geometry of rings, we will simultaneously be introducing new bond stretching parameters, whether we want them or not. (GROMOS is a notable exception, using separate atom typing for valence versus van der Waals terms [61, 84], though similar concerns still apply.) Direct chemical perception allows us to easily avoid this and focus on the (potentially unique) chemical effects which are important for individual force terms. DCP also allows the issue of generality versus accuracy to be explored specifically for individual parameters in the force field without requiring coupling among all parameters. For example, with DCP, one can easily explore whether introduction of a new Lennard-Jones parameter improves agreement with specific data, without necessarily requiring new torsions to be introduced to the force field.

Direct chemical perception also opens up new avenues for inquiry which we believe will have later payoffs for force field accuracy, simply because if parameters are assigned by processing molecules, they can be assigned using new rules. One specific example of this is the use of partial bond orders (see Section 4.4) for assigning parameters. As part of a procedure of calculating partial charges for atoms in a molecule, one can calculate Wiberg [85] bond orders and potentially use these to interpolate bonded parameters in a molecule-specific way. This procedure could capture the effects of neighboring electron-withdrawing or electron-donating groups on bond order without a need for additional force field parameters. Additionally, in our view, DCP makes force fields more easily extensible simply because the chemical perception is not hard-coded into a piece of software by an expert.

Here, we introduce one particular approach to direct chemical perception and an associated prototype small molecule force field.

### 3.3 We implement direct chemical perception to simplify force field development

In this work, we introduce a specific implementation of direct chemical perception, based on the chemical query language SMARTS [86] and its extension in SMIRKS, use it to develop a new, SMIRKS Native Open Force Field (SMIRNOFF) format, and implement an AMBER-family force field covering alkanes, ethers and alcohols in this format. We show how SMIRKS, and the SMIRNOFF format, can dramatically reduce the complexity (in terms of number of apparently independent parameters) in existing force fields while still yielding the same energies, while at the same time allowing a variety of new innovations which would be quite difficult in typical force fields. We also introduce a new force field, SMIRNOFF99Frosst, which is a prototype general small molecule force field in the SMIRNOFF format, and an AMBER-family descendant of Merck’s parm@Frosst force field [87]. We show that SMIRNOFF99Frosst covers comparable chemical space to GAFF/GAFF2 with roughly comparable accuracy.

SMARTS patterns have been used before in assigning force field parameters. For example, the Foyer effort [88-90] provides a language to encode existing force fields using SMARTS, and OpenBabel [91] uses SMARTS to encode UFF and GAFF force fields. They have also found important applications for torsion libraries and torsion angle preferences [92-94]. Richard Dixon’s OEAntechamber effort provided a proof-of-principle showing that GAFF atom typing could be encoded with SMARTS patterns [95]. One key distinction here, however, is our focus on using direct chemical perception — that is, rather than just codifying atom typing with SMARTS/SMIRKS, we assign parameters directly based on processing the molecular graph using such substructure queries.

## 4 Methodology

In this work, we introduce the SMIRKS native open force field format (SMIRNOFF), version 0.1, as an example of direct chemical perception and a way of encoding current and new force fields.

### 4.1 SMIRNOFF implements direct chemical perception using SMIRKS

#### 4.1.1 We use SMIRKS patterns for direct chemical perception

Daylight’s SMARTS patterns [86] (http://www.daylight.com/dayhtml/doc/theory/theory.smarts.html) provide a language allowing chemical substructure queries using rules that build upon the familiar Simplified Molecular Input Line Entry Specification (SMILES) format (Figure 5 and Table 1). Essentially, SMARTS allows molecules to be processed with specific substructure queries to recognize specific chemistry and provides one approach to direct chemical perception. Here, to make it easier to refer to specific atoms individually by labels, we adopt the closely related SMIRKS language [96] (http://www.daylight.com/dayhtml/doc/theory/theory.smirks.html) which was designed to handle reactions. For our purposes, however, SMIRKS serve like SMARTS patterns with numerical labels assigned to atoms (Figure 5). Thus we use SMIRKS patterns for chemical perception to determine where to utilize particular force field parameters, then process the specific molecule to finish assigning those parameters (Figure 6).

**Figure 5.**
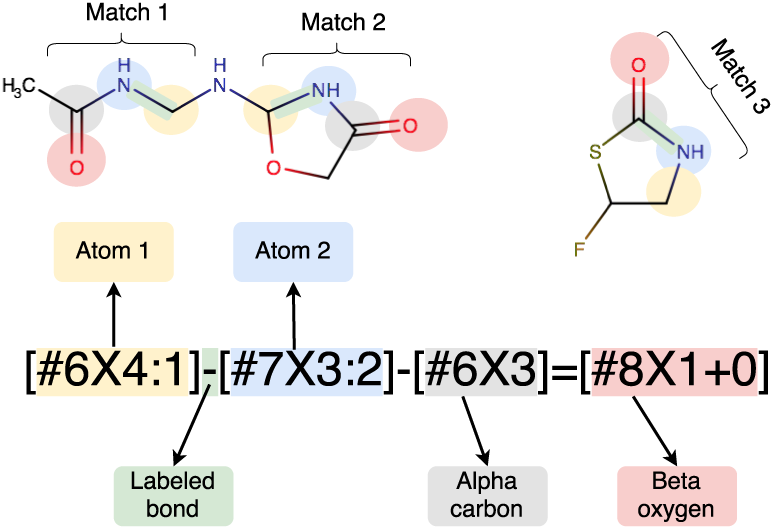
Using the SMIRKS chemical query language for direct chemical perception. SMIRKS allows for direct chemical perception via chemical substructure searches; here, a single SMIRKS pattern recognizes three different substructures in two different molecules (Match 1, 2, and 3), with matches shown by color coding of the atoms/SMIRKS patterns involved. Here, the relevant pattern is a carbon with four connections (`[#6X4:1]` (yellow) single bonded (−) to a trivalent nitrogen (`[#7X3:2]`, blue) which is single-bonded to a trivalent carbon (`[#6X3]`, gray) which itself is double bonded (=) to a neutral oxygen with a single connected atom `(#8X1+0]`, red). The first carbon and nitrogen are singled out for special treatment by having numerical atom labels (:1 and :2) assigned to them, in this case because the SMIRKS pattern would be used to assign a bond parameter to the bond connecting the two labeled atoms. The bond connecting these labeled atoms (green) is singled out for special treatment because it connects two labeled atoms. Specifically, the force field format we introduce here allows us to use this SMIRKS pattern to refer specifically to parameters for the bond connecting these two labeled atoms.

**Table 1.**
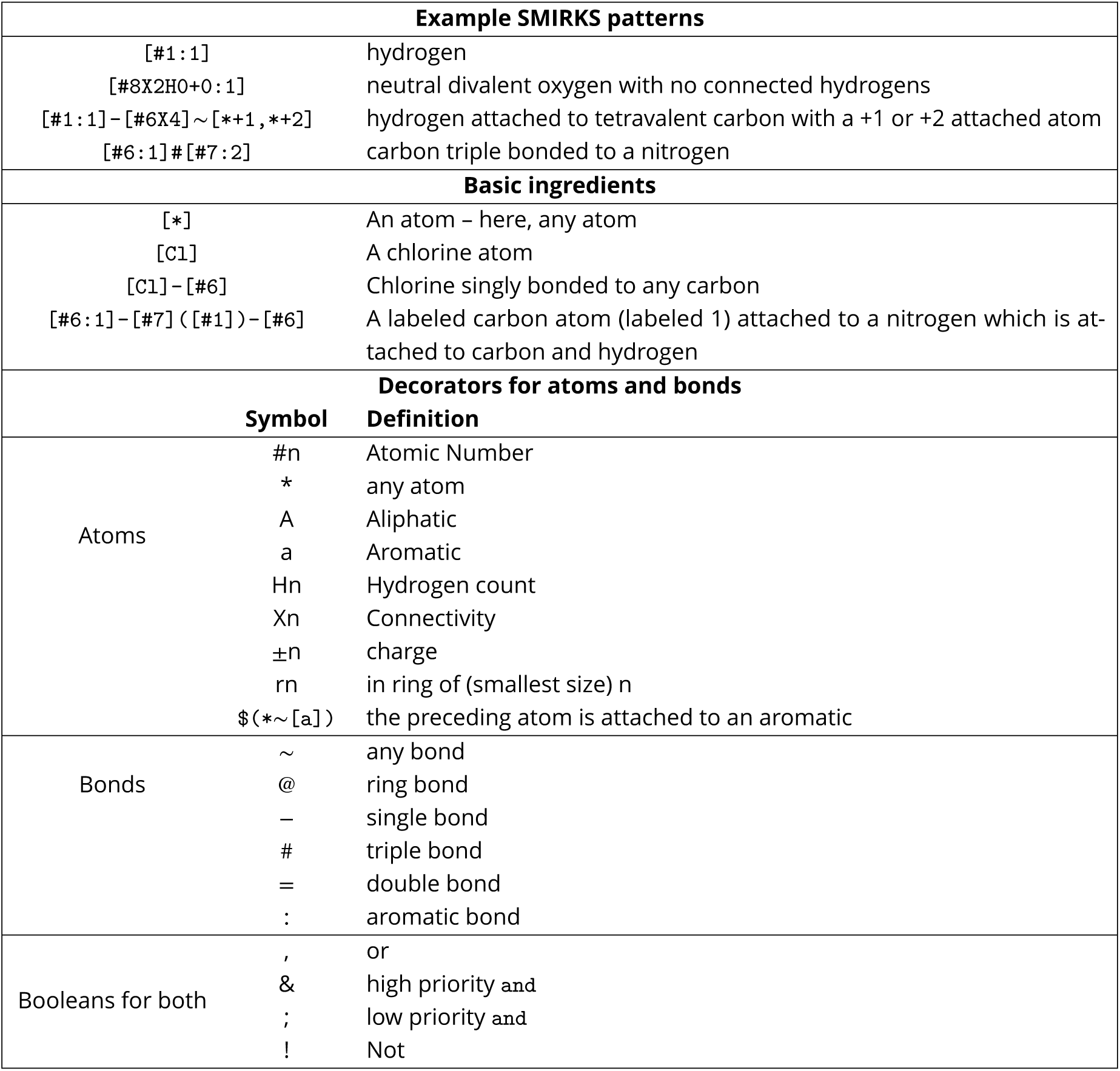
SMIRKS patterns and their ingredients. Shown are some example SMIRKS patterns (top) used in SMIRNOFF force fields, as well as selected basic building blocks of SMIRKS patterns and selected decorators used in describing atoms and bonds. SMARTS patterns are a modification of SMILES for substructure queries, such that every SMILES is a valid SMARTS pattern but not every SMARTS pattern is a valid SMILES. SMIRKS, for our purposes here, are SMARTS patterns that use numerical atom labels to refer to specific atoms when needed to allow a later reference to an atom by number, such as in `[#8X2H0+0:1]` where the atom is labeled 1 via: `1` for later reference. SMARTS/SMIRKS patterns can refer to atoms via both element numbers as well as element symbols (though it is important to note that, since it is a subset of SMILES, both `c` and `C` refer to carbon atoms, but the former to aromatic and the latter to aliphatic, and similarly for certain other elements); here we most frequently use element numbers. The OpenSmiles specification (http://opensmiles.org/opensmiles.html) provides a more complete description of SMILES patterns and the operators they can use which can supplement this table.

#### 4.1.2 The SMIRNOFF format uses SMIRKS to designate force field parameters

Our SMIRNOFF format is an XML-based format which allows specification of SMIRKS patterns and associated (unit-bearing) parameters for individual force terms in a force field, so each force term (bonds, angles, torsions, nonbonded interactions, charges, constraints, etc.) can use its own chemical perception as needed (Figure 6, Tables 2-3). The format is hierarchical and last-one-wins, so that if multiple SMIRKS substructure queries match a particular needed force term, such as a bond, the last matching set of parameters is the set actually applied. This allows easy coverage of large swaths of chemical space by first introducing general parameters and then overriding these in special cases where more sophistication is needed. Here we discuss SMIRNOFF format v0.1; subsequent revisions are pending. Most details of the format are provided in Tables 2-3); however, it is worth noting that the overall `SMIRNOFF` tag takes an optional `aromaticity_model` attribute which may be any of OpenEye’s supported aromaticity models (see Section 5.2).

**Figure 6.**
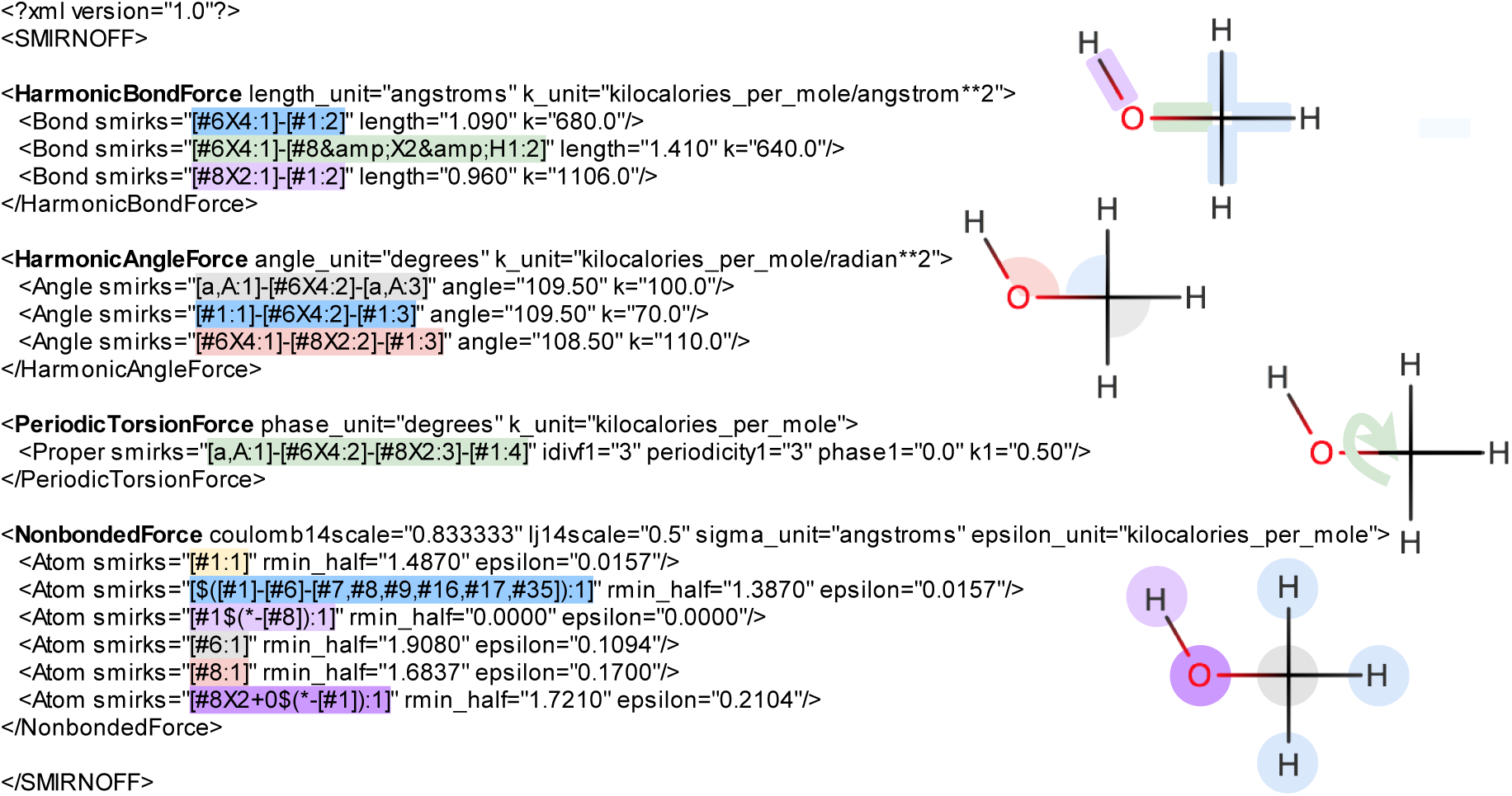
Application of a SMIRNOFF to methanol. Shown at left is an excerpt of the SMIRNOFF force field format (v0.1) XML for a test force field for our AlkEthOH test set, and the representations of methanol at right illustrate how SMIRNOFF provides the necessary force field parameters for methanol. For each force type or section in the XML (boldface), SMIRNOFF loops over the needed force terms for the molecule, and finds the last (most specialized) SMIRKS match in the XML, applying the parameters indicated there to the relevant force term. SMIRKS patterns are color coded, as are instances of the applied parameters in methanol. Force types run in sections from top to bottom, and for each of the relevant blocks (bond, angle, torsion, and nonbonded forces) here, a separate diagram is shown. For example, in the `HarmonicBondForce` section at top, for the O-H bond in methanol, the SMIRKS pattern `[#8X2:1-[#1:2]` (pink) matches and thus is selected and applied (pink highlighted bond, top right). Parameter assignments are color-coded by the element(s) involved - by the primary or central atom in the case of `NonbondedForce` and `HarmonicAngleForce` parameters (except when there is redundancy, in which case the second occurrence gets a color associated with non-central atoms), and by the central two atoms in the case of `HarmonicBondForce` and `PeriodicTorsionForce` parameters. Gray is used for carbon, red for oxygen, light green for oxygen-carbon, yellow for hydrogen, pink for hydrogen-oxygen, and light blue for hydrogen-carbon. Parameterization of some symmetry-equivalent angles is omitted in this diagram for simplicity. The hierarchical nature of parameterization is also shown here; for example, the `NonbondedForce` section contains a generic hydrogen SMIRKS pattern (`[#1:1]`, yellow) but this is overridden in the case of the hydroxyl hydrogen by a more specialized pattern (soft pink). Likewise, the generic oxygen ([`#8:1`], red) is overridden by the more specialized neutral hydroxyl oxygen SMIRKS (magenta). Not shown here are sections for bond charge corrections (BCCs) and constraints, though specifications for these are provided at https://github.com/open-forcefield-group/openforcefield/blob/master/The-SMIRNOFF-force-field-format.md. It is also worth noting that the XML format is unit-bearing, allowing handling of units utilized by various different force fields.

**Figure 7.**
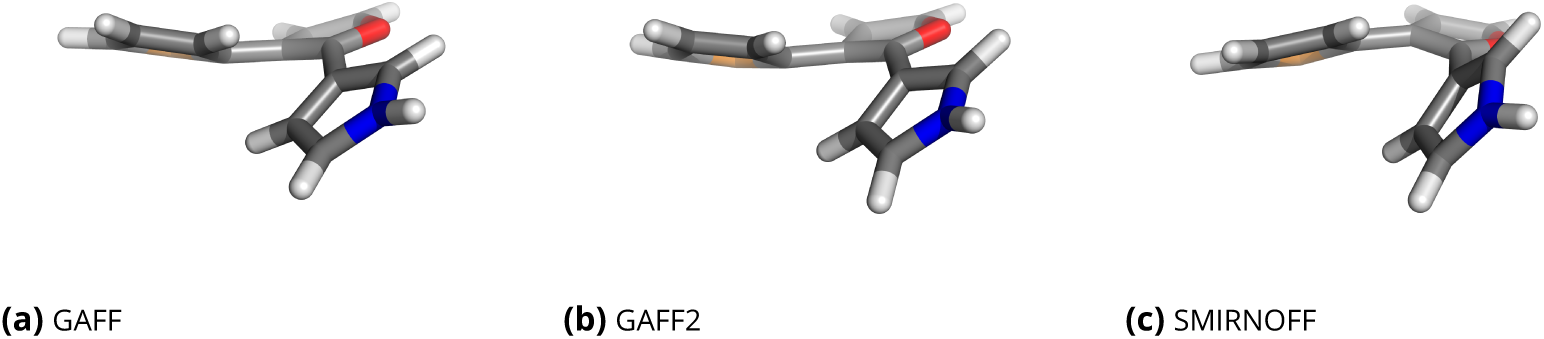
Representative geometries for the bridgehead problem case of Figure 2e. In GAFF and GAFF2, the bonds connecting the five-membered aromatic rings in (a) and (b) are non-rotatable, inducing a steric clash between protons which forces the rings to partially buckle or flex (top left of (a) and (b)), whereas in SMIRNOFF99Frosst (c) the bonds are rotatable and no buckling occurs, but instead a slight additional tilt of the rings (e.g. top left of (c)). Shown are representative geometries from short gas-phase simulations with each force field. Here, both rings have approximately the same orientation in (a)-(c), but as we show in the SI, SMIRNOFF99Frosst actually allows both rings to flip between rotamers, while GAFF and GAFF2 maintain the geometry shown in (a) and (b) throughout our simulations due to the high torsional barrier.

**Table 2.** SMIRNOFF format (v0.1) sections for nonbonded interactions. Listed are sections (force types) and the corresponding nodes and attributes for nonbonded forces in version 0.1 of the SMIRNOFF format. Each section has overall attributes (which are unit-bearing and modify the behavior of the whole section) as well as individual entries which provide details of interactions or other details of the force field. Of the sections listed here, only `NonbondedForce` entries are required, but if a section is present, all required entries must be present. Full details of the format are available online at https://github.com/open-forcefield-group/openforcefield/blob/master/The-SMIRNOFF-force-field-format.md; Figure 6 shows an example of how it is used.

| NonbondedForce |  |  |
| --- | --- | --- |
| Section attributes |  |  |
| attribute | description | required? |
| coulomb14scale | Coulomb scale factor for 1-4 interactions | yes |
| lj14scale | LJ scale factor for 1-4 interactions | yes |
| sigma_unit | Units for sigma or rmin_half | yes |
| epsilon_unit | Units for epsilon or rmin_half | yes |
| Section nodes are Atom entries |  |  |
| attribute | description | required? |
| smirks | SMIRKS pattern with one indexed atom | yes |
| rmin_half | LJ distance parameter as $r_{min}/2$ | yes, or $\sigma$ |
| sigma | LJ distance parameter as $\sigma$ | yes, or $r_{min}/2$ |
| epsilon | LJ interaction parameter as $\epsilon$ | yes |
| GBSAForce |  |  |
| Section attributes |  |  |
| attribute | description | required? |
| gb_model | Selected GB model | yes |
| solvent_dielectric | dielectric constant for solvent | yes |
| solute_dielectric | internal dielectric for solutes | yes |
| radius_units | Units for radii | yes |
| sa_model | surface area model | yes |
| surface_area_penalty | unit-bearing value for surface area penalty | yes |
| solvent_radius | solvent radius with units | yes |
| Section nodes are Atom entries |  |  |
| attribute | description | required? |
| smirks | SMIRKS pattern with one indexed atom | yes |
| radius | GB radius | yes |
| scale | GB scale parameter | yes |
| BondChargeCorrections |  |  |
| Section attributes |  |  |
| attribute | description | required? |
| method | Base charging method, e.g. AM1 | yes |
| increment_unit | units used for bond charge corrections | yes |
| Section nodes are BondChargeCorrection entries |  |  |
| attribute | description | required? |
| smirks | SMIRKS pattern with two indexed atoms | yes |
| increment | increment to move from first indexed atom to second | yes |
| Constraints |  |  |
| Section attributes |  |  |
| None |  |  |
| Section nodes are Constraint entries |  |  |
| attribute | description | required? |
| smirks | SMIRKS pattern with two indexed atoms to constrain | yes |

**Table 3.**
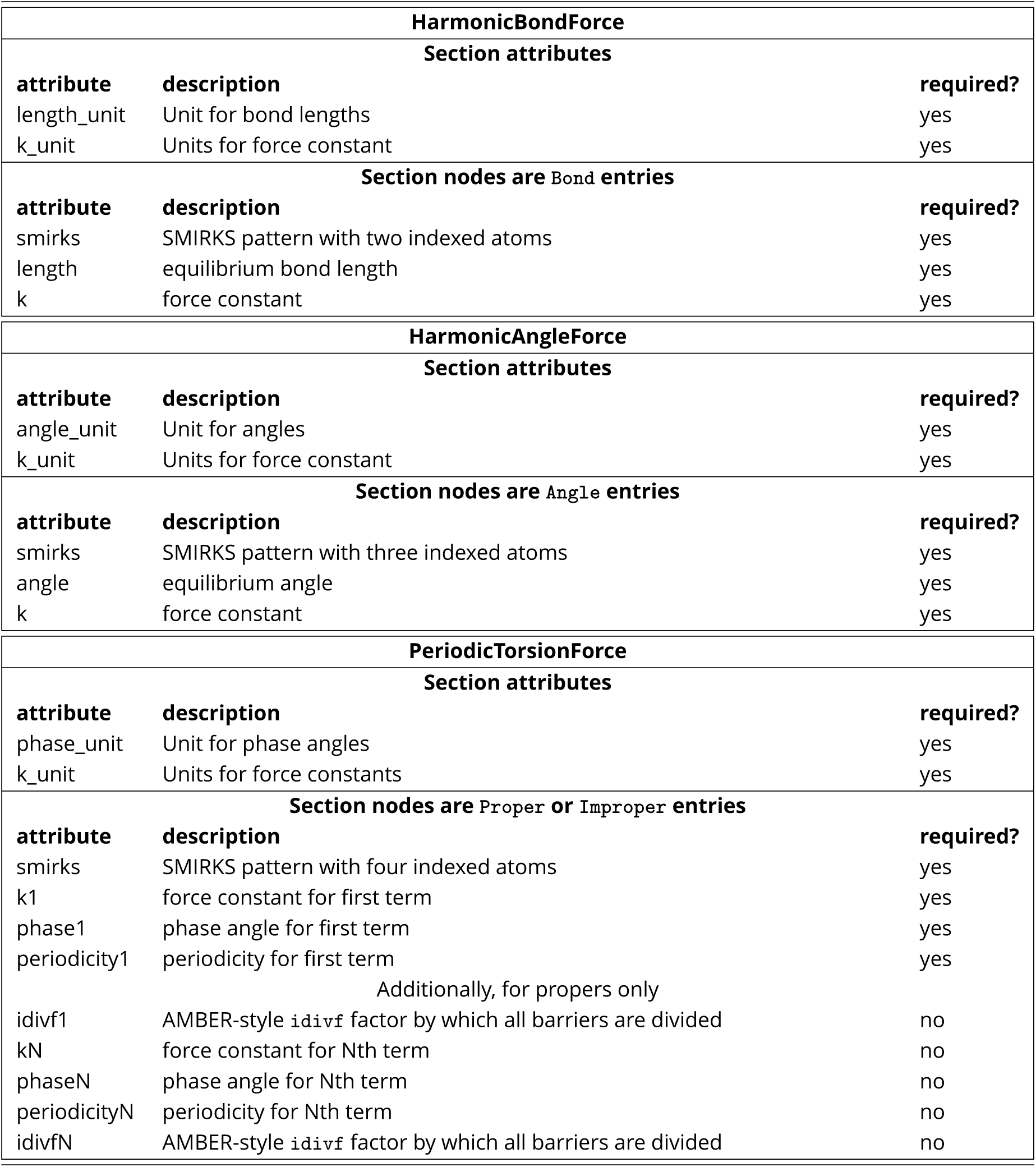
SMIRNOFF format (v0.1) sections for bonded interactions. Listed are sections (force types) and the corresponding nodes and attributes for bonded forces in version 0.1 of the SMIRNOFF format. Each section has overall attributes (which are unit-bearing and modify the behavior of the whole section) as well as individual entries which provide details of interactions or other details of teh force field. Of the sections listed here, all are required. Each section must have all required attributes; optional attributes can be omitted. Full details of the format are available online at https://github.com/open-forcefield-group/openforcefield/blob/master/The-SMIRNOFF-force-field-format.md; Figure 6 shows an example of how it is used. AMBER functional forms define the force constant *k* in a manner that differs by a factor of two from some conventions; here we do not follow AMBER conventions and elect to use the standard harmonic definition 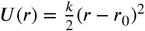. We also differ from AMBER in handling of improper torsions; AMBER picks one particular path through the torsion (which is dependent on atom ordering) and applies a single improper torsion; we take all six paths through the trefoil of the improper and apply all six impropers, after dividing the barrier height by six. For impropers, the second labeled atom is treated as the central atom.

| HarmonicBondForce |  |  |
| --- | --- | --- |
| Section attributes |  |  |
| attribute | description | required? |
| length_unit | Unit for bond lengths | yes |
| k_unit | Units for force constant | yes |
| Section nodes are Bond entries |  |  |
| attribute | description | required? |
| smirks | SMIRKS pattern with two indexed atoms | yes |
| length | equilibrium bond length | yes |
| k | force constant | yes |
| HarmonicAngleForce |  |  |
| Section attributes |  |  |
| attribute | description | required? |
| angle_unit | Unit for angles | yes |
| k_unit | Units for force constant | yes |
| Section nodes are Angle entries |  |  |
| attribute | description | required? |
| smirks | SMIRKS pattern with three indexed atoms | yes |
| angle | equilibrium angle | yes |
| k | force constant | yes |
| PeriodicTorsionForce |  |  |
| Section attributes |  |  |
| attribute | description | required? |
| phase_unit | Unit for phase angles | yes |
| k_unit | Units for force constants | yes |
| Section nodes are Proper or Improper entries |  |  |
| attribute | description | required? |
| smirks | SMIRKS pattern with four indexed atoms | yes |
| k1 | force constant for first term | yes |
| phase1 | phase angle for first term | yes |
| periodicity1 | periodicity for first term | yes |
| Additionally, for propers only |  |  |
| idivf1 | AMBER-style `idivf` factor by which all barriers are divided | no |
| kN | force constant for Nth term | no |
| phaseN | phase angle for Nth term | no |
| periodicityN | periodicity for Nth term | no |
| idivfN | AMBER-style `idivf` factor by which all barriers are divided | no |

We illustrate this here by showing application of a sample SMIRNOFF format (v0.1) force field for our AlkEthOH set (Section 4.5.1) to methanol in Figure 6, for the case where methanol partial charges are provided by the user. The minimal amount of force field information for methanol is quite brief and relatively human-readable in this case. We need three different bond stretching parameters; three angle-bending parameters, a single torsion (which involves any torsion passing with a central bond between tetravalent carbon to divalent oxygen, with hydrogen as the terminal atom), and three different nonbonded (Lennard-Jones) parameter types, though several more are shown for illustration. The Figure breaks out each force term separately and uses color coding to highlight how specific SMIRKS patterns match to particular substructures in methanol. An optional (omitted) `Constraints` section and tag can be used to constrain bonds specified by specific SMIRKS patterns, such as bonds involving hydrogen, and an attribute of the overall SMIRNOFF tag can be used to select the aromaticity model.

### 4.2 SMIRNOFF is implemented via OpenMM and the OpenEye tools

Our SMIRNOFF format parser is implemented in Python and sets up systems for use in OpenMM, but parameterized systems can easily be exported for use in other simulation packages via ParmEd. The format is implemented in the form of a `ForceField` class which is an extension/replacement of the normal OpenMM `ForceField` class but offering rather similar functionality aside from the dramatic difference in user input (here, taking a SMIRNOFF XML file rather than the different XML files utilized by OpenMM’s class). SMIRNOFF relies on the OpenEye toolkits [97-99] (free for academics for non-commercial use) for processing of SMIRKS patterns, substructure searches, and a variety of other key chemical tasks.

The process of parameterizing a system with SMIRNOFF is straightforward. First, a user loads one or more SMIRNOFF XML files via the `ForceField` class, then, to actually assign parameters, applies the `createSystem` function, providing it with an OpenMM `Topology` of the system and OpenEye `OEMol` objects corresponding to the molecules comprising the system. Various optional arguments allow the user to indicate whether to use provided partial charges or compute new partial charges, and select options such as cutoffs, periodic boundary conditions, etc. The output is a fully parameterized OpenMM `System`.

The SMIRNOFF `ForceField` class and related infrastructure (including various examples) are available free and open source on GitHub at http://github.com/open-forcefield-group/openforcefield, and a snapshot of the current release is available in the supporting information (SI).

### 4.3 SMIRNOFF can easily be applied to a variety of situations

In our GitHub repository, we provide examples applying SMIRNOFF (v0.1) force fields to set up (and in some cases conduct) simulations of a variety of different situations. These are found in the `examples` directory on the repository, a copy of which is archived here in the SI. Specifically, our examples include setup of a simple simulation of a small molecule in the gas phase, setup of a simulation of a small molecule in water, setup of simulation of a binary mixture, setup of a simulation of a mixed force field system where the ligand uses a SMIRNOFF force field and the protein uses an AMBER-family force field, and setup of a simulation of host-guest binding in water.

In our examples, simulation preparation is done entirely in Python except in the case of systems using mixed force fields (such as an AMBER force field for a protein) where use of AmberTools [100] is also necessary (via a Python wrapper).

### 4.4 SMIRNOFF solves problems with indirect chemical perception and enables new science

We find that SMIRNOFF easily solves the problems noted above for atom typing molecules with conjugated double bonds (Section 2.2), and with aromaticity perception in the biphenyl family. Specifically, simply assigning parameters based on whether bonds are single, aromatic, double or triple allows appropriate bond lengths and torsions to be assigned in these cases without the need to introduce complex atom types. To illustrate this, we applied GAFF, GAFF2, and our prototype SMIRNOFF99Frosst (Section 5.2) to 1,2,3,4-tetraphenylbenzene and found that, due to human-introduced logical errors in GAFF and GAFF2 (incorrect categorization of some of the complex combinations of atom types involved in torsions), the central ring buckles after running dynamics in the gas phase (Figure 1). In contrast, in SMIRNOFF99Frosst, though we have paid no particular attention to this series, the ring retains its appropriate planar conformation due to the SMIRKS pattern `[*:1]~[#6X3:2]:[#6X3:3]~[*:4]` used to assign parameters for aromatic carbon carbon bonds, which results in correct assignment of a torsion with a high barrier in such cases.

In this 1,2,3,4-tetraphenylbenzene case, GAFF and GAFF2 lack parameters for a variety of the torsions in the complex ring system, notably ca-cp-cq-cq, ca-cp-cq-cp, cp-cp-cq-ca, cq-cp-cq-ca, cp-cp-cq-cq, cp-cp-cq-cp, cq-cp-cq-cq, and cq-cp-cq-cp. The single bond ca-cp-cp-ca IS present in the force field and has a small barrier height of 0.795 kcal/mol. The parmchk2 program, which estimates missing parameters, assigns this same single-bond value to all of the missing torsions, even though all of these (in this system) occur in at least one case where they involve an aromatic bond within a ring. Thus, aromatic bonds within rings in this system end up with parameters better suited for an easily-rotatable single bond between aromatic rings — which ca-cp-cp-ca is intended for — and thus in GAFF/GAFF2 the ring ends up buckling during dynamics. Torsional distributions for this case are shown in the Supporting Information (SI).

GAFF/GAFF2 also lead to incorrect behavior for the case of the molecule of Figure 2e; as noted in Section 2.2, GAFF/GAFF2 make the bridgehead single bond between these rings be essentially non-rotatable, so the ring system prefers to remain planar but is forced to buckle by steric strain. SMIRNOFF99Frosst recognizes the bridgehead bond is a rotatable single bond, allowing it to rotate and to relieve the steric strain (Figure 7). Torsional distributions for this case are shown in the SI.

SMIRNOFF also provides a natural framework for handling of charge modification schemes such as bond charge corrections [101,102] as in AM1-BCC. While our initial SMIRNOFF99Frosst is intended to rely primarily on existing charge sets such as AM1-BCC, we implemented a mechanism to apply BCC corrections via SMIRKS patterns, which should allow easy re-derivation of alternate BCC schemes to be appropriate with other charge assignment procedures, or updates of AM1-BCC.

SMIRNOFF opens up avenues to explore new science such as use of partial bond orders for parameter interpolation (Section 3.2) based on local chemistry. The SMIRNOFF format supports the use of partialbond orders for this purpose, and assignment of bond stretching parameters via interpolation is already implemented (Figure 8).

**Figure 8.**
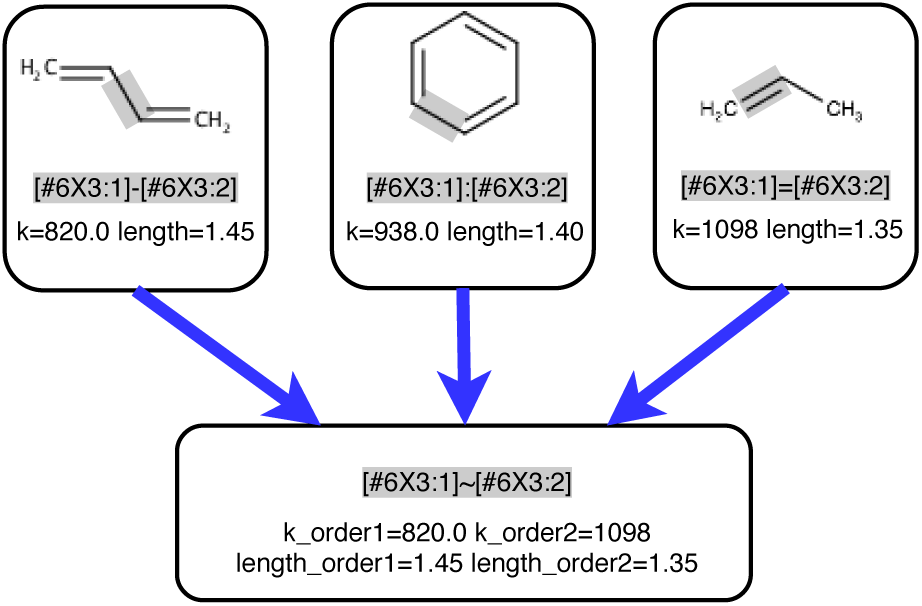
Partial bond orders can simplify the force field via interpolation. In SMIRNOFF99Frosst, there are three bond stretching parameters between carbons with three connections (top) differing in whether the bond is single, aromatic, or double. At the top we show three example molecules, with the bond in question highlighted in gray and the relevant SMIRKS expression and parameters (force constant, *k*, in kcal/(mol Å^2^), and length in Å) shown. However, an alternative approach we have implemented here is to simply use a single, more generic SMIRKS pattern `(#6X3:1]-[#6X3:2])` to recognize all bonds between carbons with three connections and to allow interpolation of parameters based on the bond order between atoms. Here, we specify a force constant and length for a bond order of 1, and a force constant and length for a bond order of 2, and with intermediate bond orders, parameters can be interpolated. For example, with linear interpolation and a Wiberg bond order of 1.5 for an aromatic bond, the force constant would be 959 kcal/(mol Å^2^) and the bond length 1.4 Å, remarkably similar to what SMIRNOFF99Frosst uses without interpolation.

SMIRNOFF also provides other opportunities to explore issues which are difficult to tackle with force fields using indirect chemical perception, such as the issue of partial nitrogen pyramidalization. That is, nitrogens can transition from being planar to being tetrahedral depending on their chemical environment (Figure 9). While existing force fields are in some cases designed to handle the planar and tetrahedral cases separately, intermediate (partially tetrahedral) cases may need to be handled as well. With SMIRNOFF, we can encode new SMIRKS patterns that recognize partially tetrahedral cases and apply appropriate parameters. For example, the following SMIRKS patterns/categorization would separate the cases considered here into planar, pyramidal, and partially planar (see Jupyter notebook in SI):

~~~
[*:1]-[#7X3:2](-[*:3])-[*:4]: pyr
[*:1]-[#7X3:2](-[#6X3:3]=O)-[*:4] : plan
[*:1]-[#7X3:2](-[#6X3:3](=[#8])-[#7])-[*:4]: part
[*:1]-[#7X3:2](-[#6a:3])-[*:4]: part
[*:1]-[#7X3:2](-[$aro_edg:3])-[*:4]: pyr
[*:1]-[#7X3:2](-[$aro_ewg:3])-[*:4]: plan
[#16$(*(=O)=O):1]-[#7X3:2](-[#6a:3])-[*:4]: pyr
[#16$(*(=O)=O):1]-[#7X3:2](-[$aro_ewg:3])-[*:4]: part
~~~

(where *plan, pyr*, and *part* denote planar, pyramidal, and partly pyramidal, respectively, `$aro_ewg` denotes an aromatic electron withdrawing group, and `$aro_edg` denotes an aromatic electron donating group). It is worth noting that in some cases such as N-methylpyrimidin-4-amine (second from right, Figure 9b), the degree of planarity depends on the conformer being considered; capturing this effect would require considerable sophistication, such as torsion terms which couple to the degree of pyramidalization — a possibility facilitated and simplified by the use of DCP.

**Figure 9.**
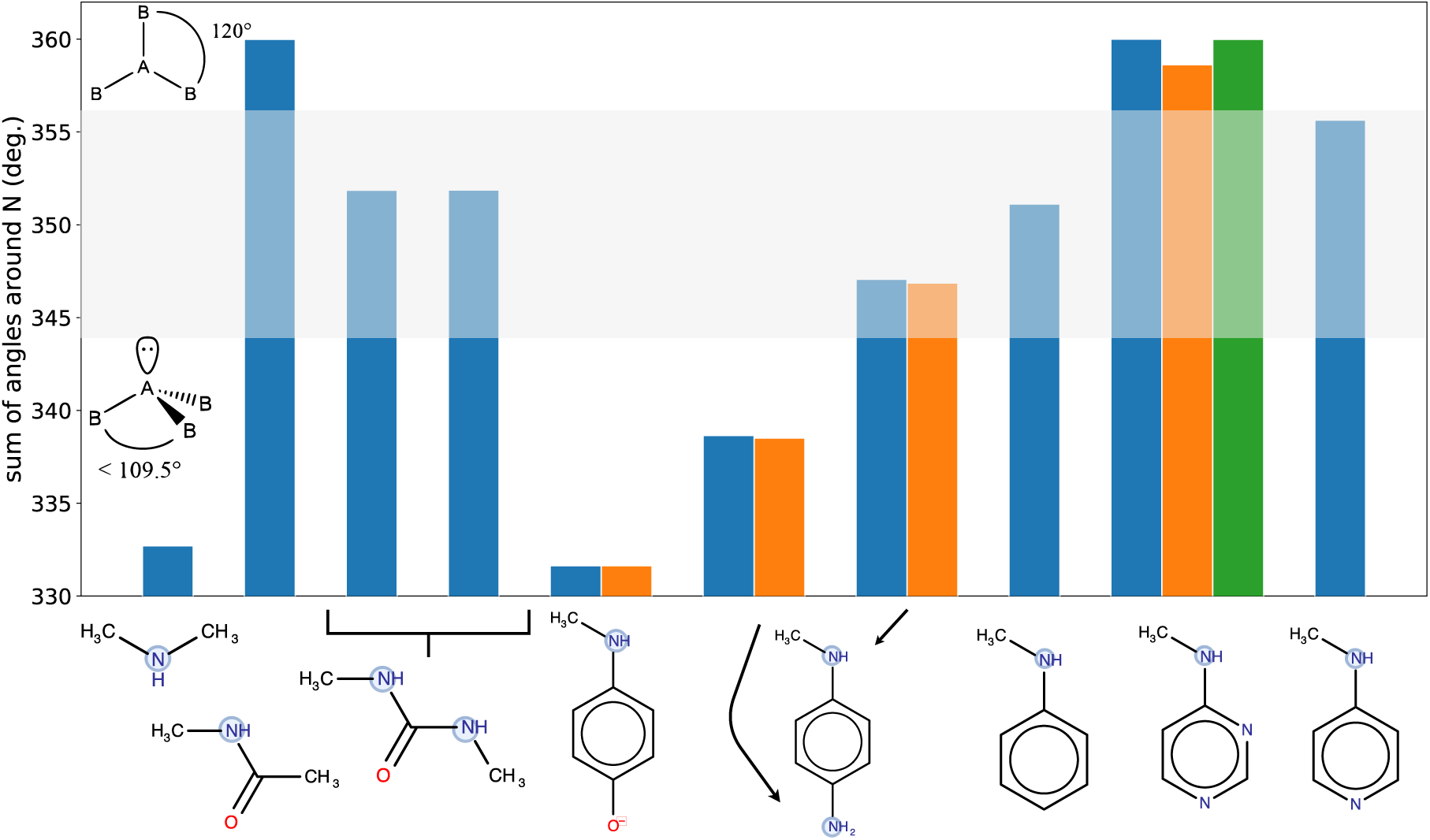
Examples of the spectrum of nitrogens showing that nitrogens can be partially pyramidal. Shown are examples of related compounds where nitrogens can transition from planar to tetrahedral (through intermediate geometries) depending on the details of their chemical environment. (A) shows 2D structures of the molecules overlaid onto a bar graph which shows, for each molecule, the sum of the three `X-N-X` angles around the nitrogen highlighted in blue (and additionally orange and green if multiple conformers have been considered). These are molecular structures after gas phase geometry optimizations with B3LYP-D3MBJ/def2-TZVP in Psi4; the sum of the angles is 360 degrees in the planar case (3*120) and 328.5 degrees in the pyramidal case (3*109.5). Partially pyramidal cases are intermediate between these values; the presence or absence of electron donating or electron withdrawing groups appears to play an important role in modulating between tetrahedral and planar. The gray band highlights the intermediate region where nitrogens are neither planar nor tetrahedral. For example, in 1,3-dimethylurea (third from left), the sum of the angles is 351.87 degrees. (B) shows the analyzed 3D structures of the molecules after gas phase geometry optimizations, in the same order as (A). In some cases, the geometries are clearly intermediate; for example, in 1,3-dimethylurea (third from left) each of the nitrogen’s protons is tilted just slightly out of the plane of the amide. Traditional atom typing requires each nitrogen geometry to have its own atom type (binning the relevant chemistry), so issues like the spectrum of nitrogen pyramidalization seen here pose challenges.

Another avenue of inquiry is the incorporation of partial charges away from atomic centers in order to better handle anisotropy, rather than maintaining exclusively atom-centered charges as used by many of our existing fixed-charge force fields. Such charges are already known to be important in a number of situations, especially for lone pairs and sigma holes, where lack of some treatment of anisotropy - either via off-site charges or via multipoles - can apparently introduce severe errors in electrostatic potentials [103] with adverse impacts on calculated properties like hydration free energies [28]. Thus, incorporation of these effects in a systematic way will likely be important for improving fixed-charge force fields, with existing work on electrostatic potentials providing guidance as to where these effects are likely most important [103]. OPLS has already moved in this direction with addition of virtual sites for halogen bonding and aryl nitrogen lone pairs in OPLS3 [9], and virtual sites have seen use in biopolymer force fields as well [1]. Virtual sites are planned to be included in an extension of the SMIRNOFF format. While off-center charges could be handled with conventional atom-type based force fields, DCP makes it considerably simpler to add them since it can be done in specific chemical environments without the need to define a wide variety of new atom types and bonded terms.

### 4.5 Methodology for validation

#### 4.5.1 Methods for energy validation versus parm@Frosst force field on AlkEthOH

To validate that force fields encoded in our SMIRNOFF format produce correct energies, we created a set of 1500 small molecules containing alkane, ether, and hydroxyl functionality in various combinations (full set available in the SI, selected molecules in Figure 12), and implemented a SMIRNOFF version of a subset of Merck’s parm@Frosst [87] force field covering this region of chemical space (`Frosst_AlkEthOH_parmAtFrosst.offxml`, available on GitHub and in the SI), then verified that we could reproduce parm@Frosst energies exactly for the entire set, as further discussed in Section 5.1.1. To validate energies, we took AMBER parameter and coordinate files produced containing parm@Frosst parameters for each molecule and loaded these into OpenMM and evaluated the energy. Then, for each molecule, we separately assigned SMIRNOFF parameters and evaluated the energy for the same conformer in OpenMM and cross-compared, verifying that the total energy and *all* of the force terms applied were exactly the same in both implementations of the force field - the original AMBER parm@Frosst implementation and our new SMIRNOFF99Frosst implementation. Code for this is available in the Supporting Information and on GitHub at https://github.com/openforcefield/openforcefield/blob/master/examples/SMIRNOFF_comparison/compare_set_energies.py, and a cross-comparison for the energy of selected molecules is included in our continuous integration testing framework which tests the SMIRNOFF implementation.

**Figure 10.**
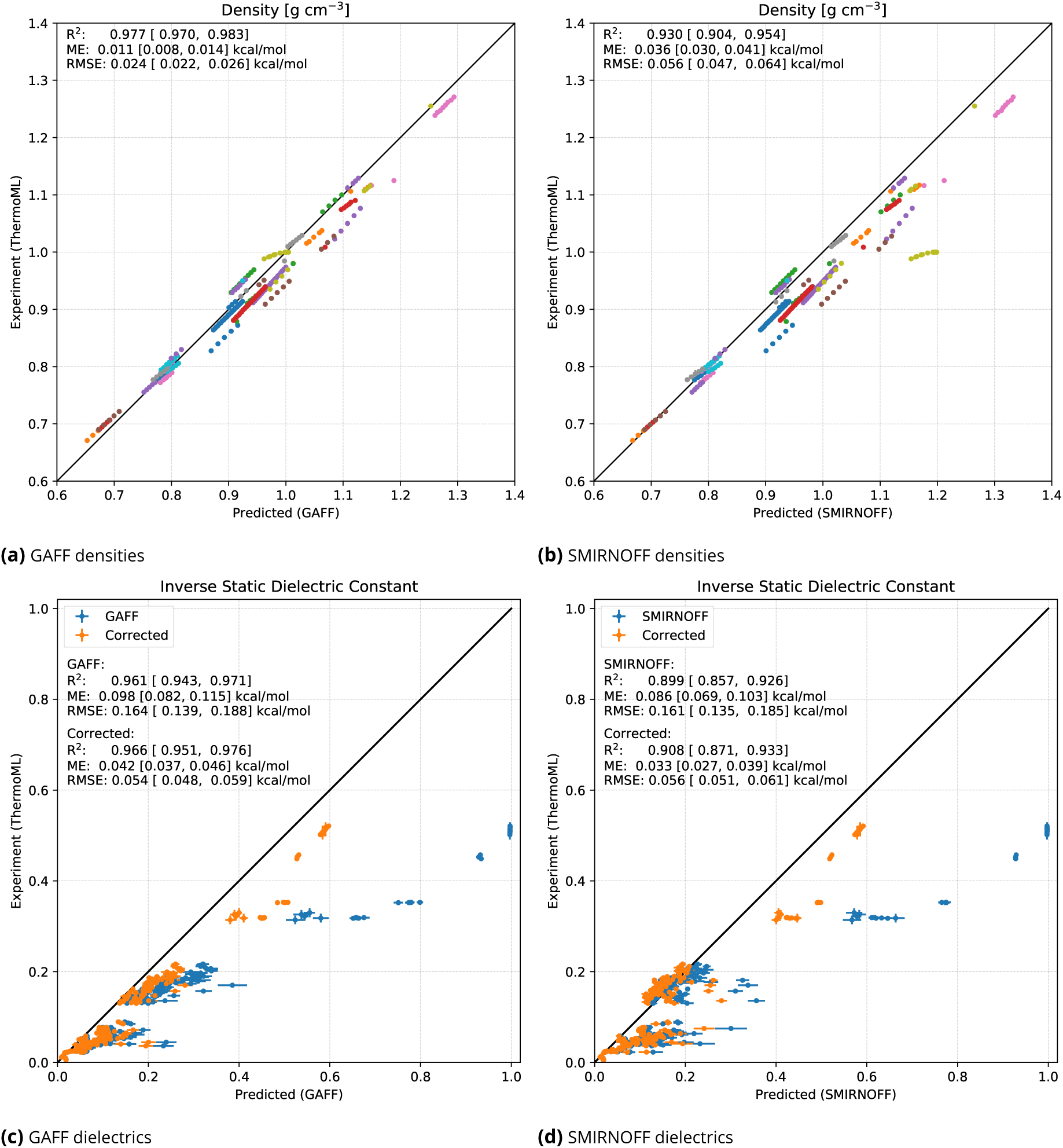
Densities and dielectric constants for pure solvents from GAFF and SMIRNOFF99Frosst. Shown are densities (top) and dielectric constants (bottom) for 45 compounds in 246 different conditions (near atmospheric pressure, at different temperatures), calculated as described in Beauchamp et al. [32] (data ours). GAFF data is shown in the left column ((a) and (c)) and SMIRNOFF99Frosst data is in the right column ((b) and (d)). In general accuracy is roughly comparable overall between the two force fields, with SMIRNOFF99Frosst densities having a slight additional systematic error relative to experiment but dielectric constants actually showing modestly improved errors (though slightly degraded correlation). Statistics are shown in the inset on each panel. In (a) and (b), each compound is shown in a different color, with multiple conditions represented in the same color. Of particular note is the density for flexible force field water, which is not typically used as a water model; in (b) this provides the most extreme set of outliers in SMIRNOFF99Frosst, around a predicted density of 1.2 g/mL and an actual density of 1.0 g/ml. A full list of compounds is available in the SI and at https://github.com/MobleyLab/SMIRNOFF_paper_code/blob/master/ThermoML_benchmark/results_GAFF/tables/data_with_metadata.csv

**Figure 11.**
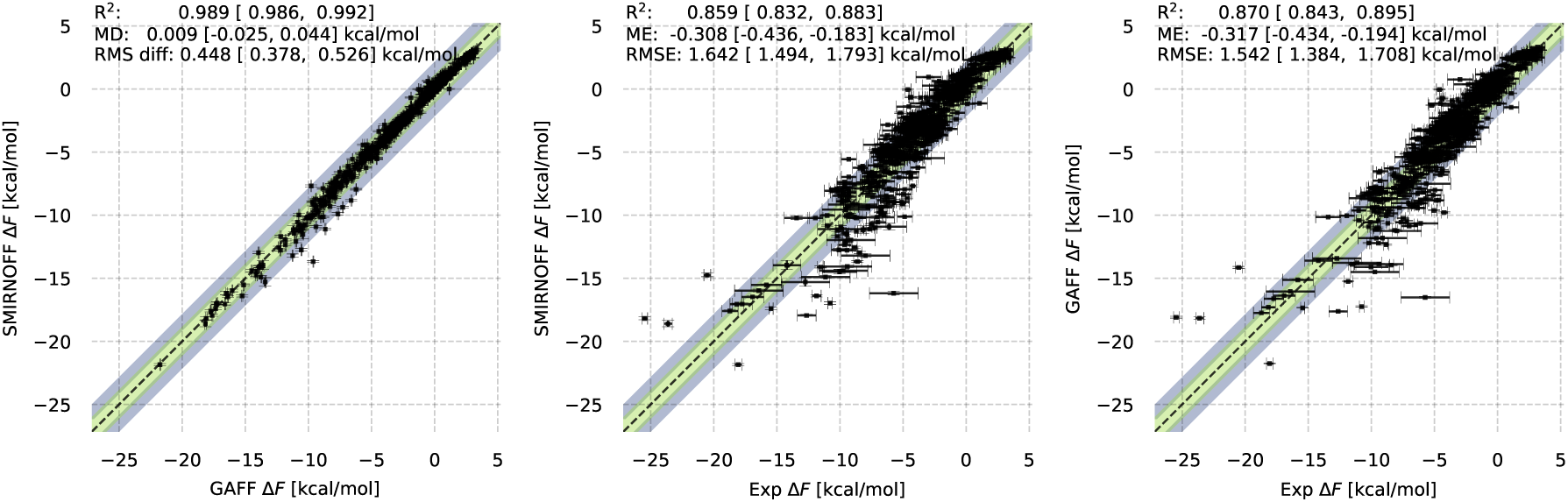
Hydration free energies for FreeSolv from GAFF and SMIRNOFF99Frosst. Shown are computed hydration free energies on the FreeSolv set for SMIRNOFF99Frosst from this work and previous work [105] with GAFF. The **left panel** shows SMIRNOFF99Frosst versus GAFF (left), the **middle panel** shows SMIRNOFF99Frosst versus experiment, and the **right panel** shows GAFF versus experiment. Statistics, with bootstrapped uncertainties representing 95% confidence intervals, are shown at the top of each panel. Here, the mean difference between SMIRNOFF99Frosst and GAFF is statistically indistinguishable from zero (left panel) though there is a significant discrepancy based on the RMS difference. However, compared to experimental values, the coefficient of determination R^2^, mean error, and RMS error for GAFF and SMIRNOFF are within confidence intervals of one another (middle and right panels) indicating that the performance of SMIRNOFF99Frosst is essentially comparable on this dataset.

**Figure 12.**
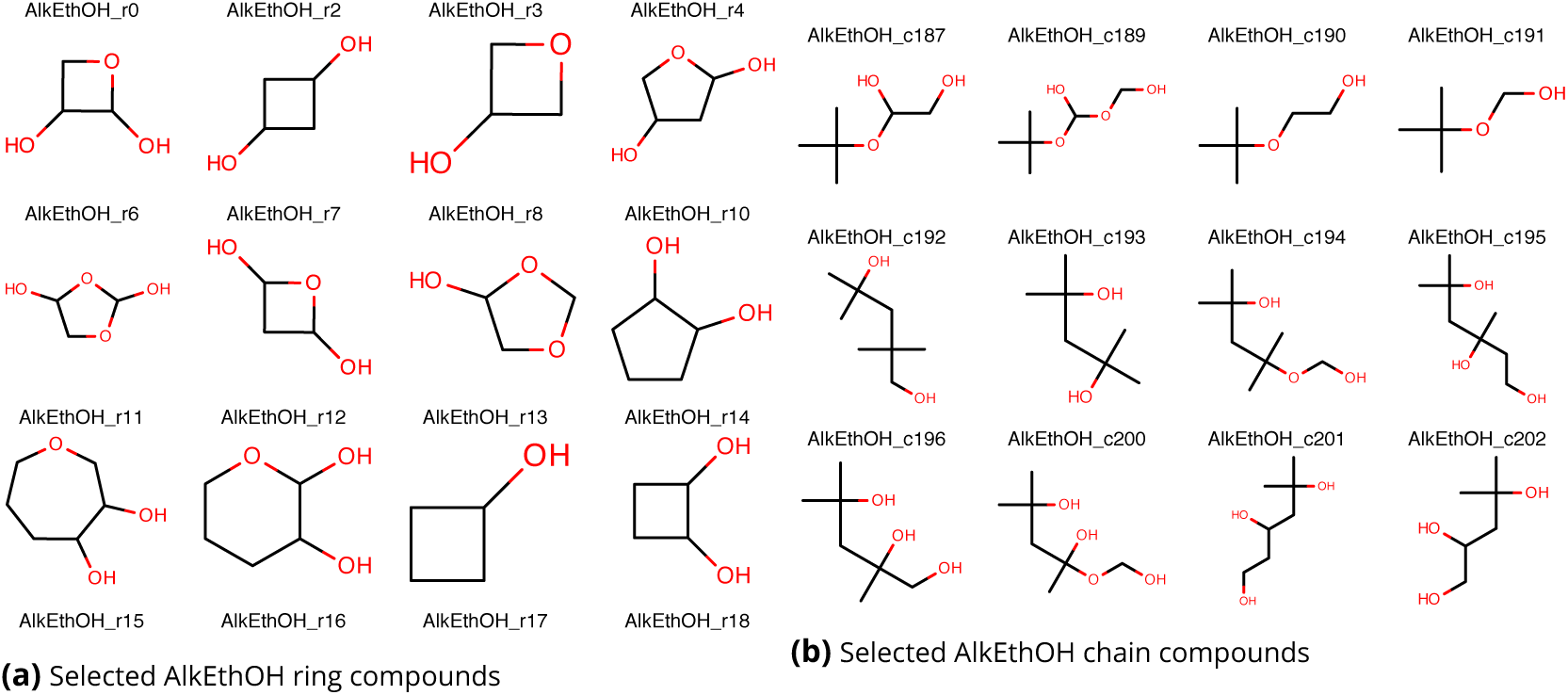
Example molecules from the AlkEthOH (alkanes, ethers, and alcohols) set. Shown are example molecules from the 1500-molecule AlkEthOH test set used here to validate energies from SMIRNOFF format against those for the Merck-Frosst force field in AMBER form at. Selected ring compounds are shown at left, and selected chain compounds are shown at right. While the set consists only of alkanes, ethers, and hydroxyls, these occur in considerable diversity. The full set of AlkEthOH compounds is provided in the Supporting Information.

#### 4.5.2 Methods for dielectric constant/density validation

To further validate the SMIRNOFF implementation on a more diverse set of molecules when our prototype SMIRNOFF99Frosst force field was available (Section 5.2), we computed densities and dielectric constants for a diverse set of small molecules previously considered by Beauchamp *et al*. [32], as further discussed in Section 5.2.6. To do this validation, we updated the pipeline of the previous work to optionally employ a SMIRNOFF force field (and put the resulting code in https://github.com/kyleabeauchamp/solutionffbench and our SI) then used it to repeat the benchmark both with GAFF and our SMIRNOFF99Frosst (version 1.0.7). All the calculations and analysis were otherwise done as described in the work of Beauchamp *et al.;* the only difference was parameter assignment; code to reproduce these calculations is deposited in the paper GitHub repository and the Supporting Information.

#### 4.5.3 Methods for hydration free energy validation

To further validate the format and our prototype force field, we computed explicit solvent hydration free energies for the FreeSolv set [104, 105] of 642 small molecules using SMIRNOFF99Frosst version 1.0.7 (see Section 5.2.6, below, for results). Hydration free energies were computed with the Yank software package [106, 107] version 0.16.0, in TIP3P [108] explicit solvent. Calculations used a temperature of 298.15 K, and a pressure of 1 atm. Calculations employed Hamiltonian replica exchange over 5000 iterations consisting each of 500 timesteps of 2 femtoseconds each. An anisotropic dispersion correction was included out to 16 Å. Exact details of our lambda protocol are given in the SI and on GitHub (https://github.com/MobleyLab/SMIRNOFF_paper_code/blob/master/FreeSolv/scripts/yank_template.yaml); we used 20 lambda values in the solution phase and five lambda values in the gas phase. Analysis was done with MBAR [109] as employed by Yank’s standard analysis framework. Full scripts for conducting and analyzing the calculations are also available on GitHub within https://github.com/MobleyLab/SMIRNOFF_paper_code.

## 5 Validation, tests, and prototypes

Here, we discuss our results of validating and initial testing of the SMIRNOFF format and several prototype force fields.

### 5.1 The SMIRNOFF format implementation was validated via several tests

#### 5.1.1 Validation on the AlkEthOH set

In Section 4.5.1, we introduced the AlkEthOH small molecule set of alkanes, ethers, and hydroxyl-containing compounds, with parameters provided by the parm@Frosst (a parm99 descendant in the AMBER family of force fields) force field [87]; to validate our format, we checked that molecule energies and forces were the same whether parameters were assigned via AMBER parameter/coordinate files or via our new SMIRNOFF format. To achieve this, CIB constructed a minimal SMIRNOFF force field by hand which reproduced the vast majority of AlkEthOH energies from parm@Frosst.

However, energies disagreed in some cases. Specifically, torsional energies differed for molecules containing atom types `H1`, `H2`, and `H3` when we compared our SMIRNOFF AlkEthOH force field and parm@Frosst. These discrepancies are apparently due to human error in construction of the parm99 force field, which includes atom types `H1`, `H2`, and `H3` for hydrogen connected oxygen bound to carbon atoms which are themselves connected to one, two, or three electron withdrawing groups, respectively. When the types `H1`, `H2`, and `H3` were introduced, the corresponding torsion `H*-CT-CT-CT` (where `H*` denotes any type of hydrogen except `HC`) was neither re-derived nor copied in from the parent atom type HC (torsion `HC-CT-CT-CT`), leaving these torsions to receive parameters for the generic `X -CT-CT-X` [69, 82]. A similar issue occurred for `H*-CT-CT-H*` for all of `H1`, `H2`, `H3`, with this torsion receiving generic parameters rather than the torsion for the parent atom type `HC` (`HC-CT-CT-HC`). An additional issue arose for `H2` and `H3`, where the `X-CT-CT-X` torsion is applied to all `H2`, `H3-CT-CT-OH`, `OS` torsions (where the comma denotes an *or*) but not to `HC`, `H1-CT-CT-OH`, `OS` torsions involving the parent atom types H1 and H2. It appears that all of these torsions were simply overlooked when the new atom types were introduced (see also Figure 13 for example molecules with these problems). Thus, these missing torsions were assigned generic torsional parameters and thus gave different energies than our SMIRNOFF implementation, which automatically treated the `H1`, `H2` and `H3` types as descendants of `HC` unless overridden. Subsequent investigation found that the same issues persist to this day in GAFF and GAFF2 13. Thus, our implementation of what parm99 was apparently intended to be (on this limited set) uncovered bugs in its implementation that persist in AMBER force fields to this day.

**Figure 13.**
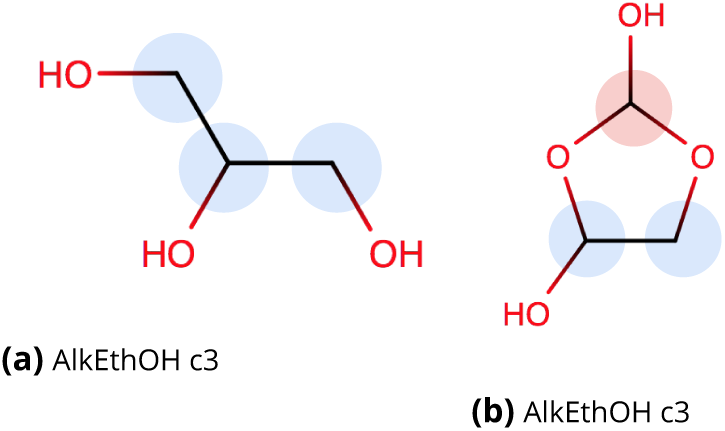
Example molecules with torsional errors in AMBER force fields uncovered in AlkEthOH. Shown are examples of molecules with torsions which were erroneously assigned generic torsional values in AMBER99, parm@Frosst, and GAFF/GAFF2 force fields, as uncovered by our testing of SMIRNOFF for AlkEthOH to ensure it could reproduce parm@Frosst energies. The errors found all involve torsions containing the `H1`, `H2`, or `H3` parm@Frosst atom types (GAFF/GAFF2 types are equivalent but lowercase), used for hydrogens connected to carbons which are themselves connected to one, two, or three electron withdrawing groups (respectively). Carbons with attached hydrogens having the `H1` atom type are highlighted in blue and carbons with hydrogens having the `H2` atom type are highlighted in red;carbons here use the `CT` atom type and oxygens use `OH` if in hydroxyls and `OS` otherwise (GAFF/GAFF2 types `c3, oh,` and `os` respectively). In (a), we had to introduce a specialized SMIRKS pattern `[#1:1]-[#6X4:2](-[#8])-[#6X4:3]-[#6X4:4]` to reproduce parm@Frosst energies on this set because parm@Frosst applies the generic `X-CT-CT-X` torsion (barrier height 1.4/9 kcal/mol) to all `H*-CT-CT-CT` torsions *except* `HC-CT-CT-CT` (parm@Frosst barrier height 0.16/3 kcal/mol). We also had to introduce a specialized SMIRKS pattern `[#1:1]-[#6X4:2]-[#6X4:3](-[#8])-[#1:4]` and associated parameters to reproduce parm@Frosst energies because parm@Frosst applies the generic `X -CT-CT-X` torsion (parm@Frosst barrier height 1.4/3 kcal/mol) to all `H*-CT-CT-H*` torsions *except* `HC-CT-CT-HC` (parm@Frosst barrier height 0.05 kcal/mol). In (b), the same two patterns also occur, but we additionally had to introduce the specialized SMIRKS pattern `[#1:1]-[#6X4:2](-[#8])(-[#8])-[#6X4:3]-[OX2:4]` to reproduce parm@Frosst energies where the `X -CT-CT-X` torsion is applied to all `H2,H3-CT-CT-OH,OS` torsions (where the comma denotes an “or”). In this case, `HC,H1-CT-OH,OS` torsions get use two terms, with a periodicity 3 barrier height of exactly zero and a periodicity 1 barrier height of 0.25 kcal/mol, whereas the generic has a single term with a periodicity 3 barrier height of 1.4/9 kcal/mol. These three problematic torsions occur for many molecules in our set, and all three issues persist in GAFF and in beta versions of GAFF2 (such as the AmberTools 16 and 17 versions examined here). These issues appear to be a case of human error, where introduction of new derivative atom types (H2 and H3 are derivatives of H1) was not accompanied by reproducing or refitting some of the relevant torsional potentials. Thus, to reproduce parm@Frosst energies we actually had to create two versions of our AlkEthOH force field - a minimal one which reproduces parm@Frosst as it was intended for these molecules, and a more extensive one which also reproduces these three bugs in parm@Frosst.

To finish validating our SMIRNOFF format, then, we extended our prototype AlkEthOH force field to deliberately apply generic torsional parameters to the chemical environments where they had been erroneously applied in parm99 and parm@Frosst. This required introducing several additional parameters and SMIRKS patterns (Figure13), making the force field *more complex* to recapitulate the bugs in the original force field. Once this was done, we perfectly reproduce parm@Frosst energies and forces for the entire AlkEthOH set. However, we believe the need to make the force field more complex to reproduce bugs in the existing force field is evidence of how the simplicity of SMIRNOFF actually opens the way to a better handling of the underlying chemistry.

#### 5.1.2 Validation of partial bond orders for parameter interpolation

The SMIRNOFF format supports the use of partial (i.e. Wiberg) bond orders for parameter interpolation as an advanced or experimental feature (see Section 4.4 and Figure 8). Currently we support only parameter interpolation for `HarmonicBondForce` terms but plan extension to angle and torsional terms. Here, we validated our implementation based on assigning bond parameters for aromatic rings. Specifically, the `HarmonicBondForce` section of the SMIRNOFF format allows an optional tag, `fractional_bondorder` which specifies a parameter interpolation scheme, with linear interpolation (`interpolate-linear`) currently the supported method. We validated our implementation by checking that the correct parameter is assigned to benzene (in this case, only a single `[#6X3:1]!#[#6X3:2]` SMIRKS pattern is needed for carbon-carbon bonds except for triple bonds) and that the resulting interpolated bond distance and force constants are correct for the case of benzene, as shown in Figure 8.

### 5.2 SMIRNOFF99Frosst: A new general small molecule force field

#### 5.2.1 SMIRNOFF99Frosst is a prototype force field constructed by hand

SMIRNOFF99Frosst was constructed by hand from the parm99 descendant parm@Frosst force field, by CIB, who was involved with the original development of parm99 [81] and parm@Frosst [87]. The basic goals of its construction were:

1. Build a good but simple starting point for future force field improvements, preferring generality and coverage over accuracy
2. Take advantage of the SMIRNOFF format to dramatically reduce the number of parameters and simplify the force field, especially when unnecessary specialization seemed to have been added
3. Capture as much of the key chemistry as possible as simply as possible
4. Remove redundancy and near redundancy (such as by combining parameters which differ only slightly) as much as possible with the idea that if additional specialization is needed, it will be added back in subsequent force fields in a data-driven manner

The choice of aromaticity model is an important part of chemical perception, and is particularly important for perceiving fragments using SMIRKS patterns. In particular, a SMIRKS pattern for an aromatic bond between two aromatic atoms can only match a bond which is determined to be “aromatic” and not a formal double or single bond. While this is desired for many kinds of aromatic six-membered rings such as benzene and pyridine, it is a liability for five-membered heteroaromatic rings such as imidazole or oxazole. In six-membered rings, such as benzene and pyridine, there is no significant difference between the formally drawn single and double bonds. For those molecules all six bonds should often be treated identically. But this is not necessarily the case for five-membered rings. Consider imidazole as a specific example: Empirical evidence shows that the bonds between carbon and nitrogen are not all identical; some have more single bond character while others more closely resemble double bonds. The Supporting Information provides more detailed analysis of both of these possibilities. If the bonds in imidazole were all perceived as aromatic, the SMIRKS required to correctly describe the differences in these valence terms would be significantly more complex, most likely incorporating all atoms in the ring. If the bonds in the five-membered rings are identified as formal single or double bonds then the SMIRKS only require information about the atoms and bonds directly involved in the valence term.

Because of the simplification possible if we treat many five-membered rings as alternating single and double bonds, we selected the MDL aromaticity model for use with SMIRNOFF99Frosst — specifically, OpenEye Scientific’s OEChem Toolkit implementation of the MDL model (here, “OpenEye MDL” for short)^1^. The OpenEye MDL model assigns aromaticity based on a limited definition of Hückel’s 4*n* + 2 rule; the only pi-electrons counted are those coming from double bonds in the ring system. This means the electrons on hetero-atoms or anions in five-membered rings are not counted as contributing to the ring system. In other words, the only atoms and bonds perceived as aromatic with this model are those in ring systems with an odd number of alternating double bonds. This includes six membered rings with alternating single bonds such as benzene or pyridine and also fused systems such as azulene (a fused seven-membered and five-membered ring system). The ForceField class (and the SMIRNOFF format) can optionally perceive aromaticity (via specification of an aromaticity model) or use a molecule as provided, but SMIRNOFF99Frosst was designed assuming the use of OpenEye’s MDL aromaticity model and specifies this in its XML. RDKit has this same OpenEye MDL aromaticity model available in versions 2017.09.3 and later, and an RDKit-based implementation of the ForceField class is under development.

#### 5.2.2 SMIRNOFF99Frosst was created by adapting parm@Frosst and parm99

To develop SMIRNOFF99Frosst, we began by creating two simplified SMIRNOFF force fields — one based on the AMBER parm99.dat parameter file and and one on the parm@Frosst parameter file — via the following procedure:

1. The parameter file was converted section by section separately, i.e. bonds separately from valence angles, torsions, etc.
2. Each connected set of AMBER atom types was transliterated into a primitive SMIRKS string using an automated search-and-replace process. Atom types were converted into the element+hybridization SMIRKS corresponding to the atom type, e.g. “[#6X4]” for AMBER atom type CT and “[#6X3]” for AMBER atom types `C,CA,CM,C*` etc. Thus the AMBER valence angle `CT-OH-HO` was transliterated as “[#6X4:1]-[#8X2:2]-[#1:3]”. The AMBER parameter was copied into the comment field for reference in the fourth stage.
3. The whole section of parameters was then lexically sorted based on the SMIRKS version of the parameter, thus grouping together parameters having identical SMIRKS representations. Because of the oversimplified SMIRKS representation of the atom types, different parameters by AMBER atom types could have the same SMIRKS representation. The lexical sort thus tended to group together parameters of similar chemistry.
4. Groups with an identical SMIRKS representation were then manually edited according to several heuristics:

a. The default bond order of 1 (carried over as the hyphen from the AMBER atom-type-based parameter) was modified as chemically required, for example the carbonyl-containing angle `CT-C-O` transliterated as “[#6X4:1]-[#6X3:2]-[#8X1]” had to be modified to “[#6X4:1]-[#6X3:2]=[#8X1]”. Resorting was required after this. The SMARTS/SMIRKS bond orders in Table 1 were used, including the wild-card bond order “~” where appropriate (especially for generic parameters in stage 4c below).
b. Where distinct parameters required it, additional chemical complexity was introduced into the SMIRKS patterns to reflect the additional chemical complexity in the AMBER atom types that had been oversimplified in stage 2. Resorting was required after this.
c. Identical parameters and highly similar parameters were collapsed to a single occurrence representing a more generic parameter. This was where the greatest reduction in the number of parameters occurred.
5. Occasionally the collapsing of redundant parameters as in stage 4c above could be generalized to include greater generality even between the element+hybridization SMIRKS of stage 2 above; this was especially true for valence angles involving an aromatic carbon as the middle atom.
6. The section of parameters was manually reordered to get the correct “last one wins” behavior wherein subsequent more specific parameters can overwrite earlier more general parameters.

The SMIRNOFF for each of parm99.dat and parm@Frosst were then combined (section by section) in a process identical to stage 6 above, removing the occasional redundancy as in stage 4c above. This produced an initial draft of SMIRNOFF99Frosst.

CIB worked from the relevant parm99 and parm@Frosst parameter sets in AMBER and AMBER `frcmod` formats, modifying these to contain SMIRKS patterns in the first column in a format we call a SMIRNOFF-ish `frcmod` file. These custom files were then converted to the XML-based SMIRNOFF format via a custom Python script, `convert_frcmod.py` available in our SI and on GitHub at https://github.com/openforcefeld/openforcefield/tree/master/utilities/convert_frosst.

Subsequent extensions of SMIRNOFF99Frosst to improve coverage of chemical space were made by CCB by modifying/extending the same format, including by generalizing and importing some parameters from GAFF2 as further detailed on GitHub at https://github.com/openforcefield/smarty/pull/232

Some of the authors’ previous work had revealed problems with the zero Lennard-Jones parameters used for hydroxyl hydrogens in AMBER-family force fields. Un some situations, these actually allow hydroxyl hydrogens to sit on top of neighboring polar atoms such as oxygen atoms in adjacent molecules or residues, resulting in crashes, as we detail in the Supporting Information (DLM and CIB, unpublished work). Thus we took this opportunity to also add very small but nonzero Lennard-Jones radii and well-depths to hydroxyl hydrogens (see Supporting Information) and included these in all of our validation work.

The resulting SMIRNOFF99Frosst force fields are available in a versioned manner on GitHub at https://github.com/openforcefield/smirnoff99Frosst.

#### 5.2.3 SMIRNOFF99Frosst covers comparable chemical space to GAFF/GAFF2 with 332 lines of parameters, 20 times fewer than GAFF/GAFF2

As discussed above, when generating parameters we attempted to cover the same general organic chemistry space that is covered by the AMBER (and GAFF) force fields, so we checked how much chemical space SMIRNOFF99Frosst covers. However, “coverage” needs to be defined carefully due to frequent use of generic parameters. In SMIRNOFF99Frosst the first parameter in each section uses a generic SMIRKS pattern which could match any parameter of that type. For example, the SMIRKS for the generic bond parameter is “`[*:1]~[*:2]`”. Given the hierarchical nature of the SMIRNOFF format, if a more specific parameter exists for a given bond (or any parameter type), it will not be assigned the generic parameter. When checking for coverage, we typed molecules with SMIRNOFF99Frosst and then checked for molecules where a generic parameter was assigned; when the generic is used, a molecule is considered “not covered” by the force field, so we would have full coverage of a set of molecules if no generic parameters are assigned. In contrast, conventional force fields (GAFF and parm@Frosst) do not cover a molecule if they fail to assign parameters. The scripts used to check for this are available on GitHub at https://github.com/openforcefeld/openforcefield/tree/master/utilities/SMIRNOFF_vs_frosst.

A series of molecule sets increasing in size and complexity were considered when evaluating coverage compared to parm@Frosst and GAFF force fields.

##### FreeSolv coverage

FreeSolv is a database of small molecules and their hydration free energies maintained by the Mobley group available at https://github.com/mobleylab/FreeSolv. All three force fields being considered, GAFF, parm@Frosst, and SMIRNOFF99Frosst, can completely parameterize the FreeSolv molecule set.

##### ZINC coverage

We also considered a subset of the ZINC database[110, 111] which was originally curated to test the parm@Frosst force field [87]. In this set SMIRNOFF99Frosst fails to cover 13 molecules out of 7505, while GAFF fails to cover 77 and parm@Frosst 3575. Five molecules in the ZINC set have inappropriate valency — that is, a carbon or nitrogen with a bond order greater than four bonds, so at least five should probably not be covered.

##### DrugBank and eMolecules coverage

Next, we moved on to DrugBank [112-115] and eMolecules (https://www.emolecules.com) which are both large databases available online for non-commercial efforts. Given the diversity covered in these sets, some molecules were filtered out before attempting to assign force field parameters. Currently SMIRNOFF99Frosst, parm@Frosst, and GAFF do not cover any metals, metaloids, or boron so we removed molecules with these atoms. We also removed molecules with greater than 200 heavy atoms or inappropriate valency. We removed an additional 66 entries in DrugBank which contained more than one molecule. Finally, we generated 3D conformations and assigned AM1-BCC charges using the OpenEye toolkits, and removed all molecules for which this process failed, resulting in 5497 and 5,718,564 molecules from DrugBank and eMolecules respectively. For DrugBank, 15 molecules were not covered by SMIRNOFF99Frosst, compared to 32 with GAFF and 2183 for parm@Frosst. For eMolecules, 30,338 molecules were not covered by SMIRNOFF99Frosst, compared to 386,891 with GAFF.

Thus, overall, we find that SMIRNOFF99Frosst covers similar chemical space to the GAFF force field (but marginally more) and substantially more than parm@Frosst.

##### Number of parameters compared to GAFF and parm@Frosst

At some level, making a simple force field which covers vast chemical space is unimpressive if it is achieved by an overly general but inaccurate force field, so accuracy, in addition to coverage and simplicity, is a key criteria as we address in Section 5.2.6; we find the accuracy of SMIRNOFF99Frosst is roughly comparable to GAFF. Yet substantial coverage is achieved with comparatively few parameters, in part because of elimination of redundancy and additional parameters introduced by atom typing, and in part because of design intent (our preference for a simple starting point). Thus, SMIRNOFF99Frosst currently has only 332 lines in the full force field XML file in version 0.1 of the specification, counting headers and force type labels. The number of actual parameters present varies somewhat across force types with bond, angle, and nonbonded terms having two parameters per line and torsional terms having two or more parameters per line. This is in contrast to parm@Frosst with 720 lines of parameters from parm99 and another 2893 from the parm@Frosst extensions [87] for a total of 3613 lines of parameters, GAFF with more than 6300 lines of parameters, and GAFF2 with more than 6700 lines of parameters. Thus, SMIRNOFF99Frosst would be a simpler starting point for automated parameter fitting machinery such as ForceBalance [52-56] because dramatically fewer parameters would need fitting. It is likely that additional parameters will need to be introduced (via refinement of existing parameters or splitting out some new parameters) to make accuracy comparable to GAFF, GAFF2, or other force fields such as OPLS2005 [116] or OPLS3 [9], as we will explore in future work, but we are confident that SMIRNOFF’s approach to chemical perception will allow the number of parameters to stay relatively lower.

#### 5.2.4 SMIRNOFF99Frosst provides a starting point for further development

In our view, then, SMIRNOFF99Frosst will be a good starting point for future force field development. The relatively simple chemical perception employed and the hierarchical format mean that extension to more sophisticated chemistry is straightforward and can be done by introducing new SMIRKS patterns further down in the hierarchy, where needed, to cover special cases. It will be interesting to experiment with refitting all of the parameters given the chemical perception codified in this prototype force field and to see how much these change, and additionally to explore refitting only where chemistry dictates it is needed. For example, we could generate quantum mechanical reference data for a wide range of molecules and use SMIRKS queries to identify groups where a specific parameter is used, for example for a bond length. When the distribution of bond lengths from the reference data for a specific SMIRKS query is multimodal, it may suggest that additional parameters need to be introduced. Our longer-term goal is to automatically sample over chemical perception *while* fitting the force field in order to let data (rather than human expertise) drive what chemical perception is employed by the force field. Parallel work on SMIRKY [117], our tool for sampling chemical perception, will facilitate this. Specifically, SMIRKY, which uses Monte Carlo sampling to explore chemical perception space (in this case represented by SMIRKS patterns) couples with this work as first steps towards our larger vision. We want to change chemical perception *while* parameterizing force fields, allowing data to drive both the selection of which parameters are present and what values they have, fitting or sampling over both chemical perception and parameters simultaneously.

#### 5.2.5 SMIRNOFF99Frosst is relatively simple

SMIRNOFF99Frosst seeks to encode the minimum amount of chemical complexity necessary to have a general-purpose small molecule force field for organics. For example, sp^2^ carbon single bonds have just a single bonded parameter line. Double bonds between sp^2^ carbons likewise have only a single bonded parameter. Angle parameters are as condensed as possible, in many cases depending only on the chemistry of the central atom. Additionally, for angles, relatively little attention has been paid to angle parameters within rings, with the assumption that ring geometry will tend to be dictated (to first order) based on topological constraints rather than specifics of angle parameters. Torsions exhibit considerable additional complexity, but barrier heights are estimates derived from the original force field while attempting to simplify parameters without introducing significant errors. As each new element of chemical perception (each SMIRKS pattern in this case) essentially bins chemistry to some extent, the goal here was to cover all conjugation classes and as much chemistry as possible with few bins. Thus, SMIRNOFF99Frosst has widespread use of relatively generic SMIRKS patterns, but we hope it captures much of the essential chemistry.

#### 5.2.6 We validated SMIRNOFF99Frosst by calculating dielectric constants, densities, and hydration free energies

To test our SMIRNOFF99Frosst prototype force field, we performed an extensive benchmark on density and dielectric constant calculations of pure solvents, as well as an even larger test on the FreeSolv database of hydration free energies, comparing SMIRNOFF99Frosst results with those from GAFF.

As discussed in Section 4.5.2, density/dielectric constant calculations used the test set from prior work [32] with minor updates to the code; we repeated all calculations with both GAFF and SMIRNOFF99Frosst to ensure the only difference was the force field. Results are shown in Figure 10. We find that the accuracy of calculated densities and dielectric constants is quite similar to GAFF, with densities having a marginally larger systematic error than GAFF in the direction of being too high relative to experiment, but dielectric constants actually performing marginally better than GAFF relative to experiment in terms of average error. On the whole performance is roughly comparable, and suggests that SMIRNOFF99Frosst is indeed a good starting point for further development, especially given its relative simplicity.

FreeSolv hydration free energies provided an even more extensive benchmark, with 642 different values in the test set. Method details are given in Section 4.5.3 but calculations used explicit solvent alchemical free energy calculations with a similar protocol (though a different software package) than previously published work with GAFF [105], allowing direct comparison of calculated values such that the primary differences should be force field differences. Figure 11 shows the results of this comparison, with (a) showing comparison to previously published values, (b) showing comparison of SMIRNOFF99Frosst results with experiment, and (c) showing comparison of prior GAFF results with experiment. Overall, SMIRNOFF99Frosst and GAFF agree extremely well, with a Pearson R of 0.989 (95% confidence interval (CI) [0.986, 0.992]), mean difference of 0.009 [-0.025,0.044] kcal/mol, and RMS difference of 0.448 [0.378,0.526] kcal/mol. Very slight differences in performance relative to experiment are observed, but these are within confidence intervals and depend on the metric examined (e.g. the mean error is smaller with SMIRNOFF99Frosst but the RMS error is smaller with GAFF). On the whole, this further validates the notion that SMIRNOFF99Frosst has roughly comparable accuracy to GAFF despite its relative simplicity.

It is important to note that our validation work primarily assesses bulk properties and nonbonded terms, which is critically important but does not provide a great deal of insight into the accuracy of the valence terms with respect to geometries and energies. This will be tested in subsequent work and will be a major part of improving SMIRNOFF99Frosst and building subsequent force fields.

### 5.3 SMIRNOFF force fields can be used in most MD codes

The `ForceField` class used to interpret the SMIRNOFF v0.1 XML files provides essentially a drop-in replacement for the OpenMM `ForceField` class utilizing modified inputs, allowing easy application to a wide variety of different types and easy export for use in most major simulation packages. In OpenMM, use currently requires a (free for academics) license to the OpenEye Python toolkits, though an RDKit-based implementation is in progress as well. The `ForceField` class first loads and parses a force field, and then the `createSystem` function can apply that force field to a specific system. As input, it requires an OpenMM `Topology` describing the system to be parameterized, as well as a set of OpenEye `OEMol` objects for the individual molecules comprising the system, and various arguments concerning charging method, cutoff, and other choices for the system to be set up. The final result is an OpenMM `System` object containing all the information on how to compute energies and forces which then can be used for molecular simulation or modeling applications. This `System` object can be readily converted for use in other codes via ParmEd (http://parmed.github.io/ParmEd) or InterMol (http://intermol.readthedocs.io/en/latest/), allowing export to GROMACS, AMBER, DESMOND, CHARMM, and LAMMPS formats [118] for use in a wide variety of packages.

Our long-term goal is to switch to direct chemical perception for both biopolymers (proteins and nucleic acids) and small molecules, providing a natural and consistent way to handle nonnatural amino acids, posttranslational modifications, and covalent inhibitors along with ligands, cofactors, and biopolymers. At present, however, SMIRNOFF is mainly oriented towards handling of small molecules rather than biopolymers, primarily because we have not yet handled fragmentation schemes for charge assignment for consistent handling of large molecules consisting of repeating units. Thus, in the interim it may be desirable to use SMIRNOFF force fields in a similar manner to how GAFF and GAFF2 are used—to parameterize solutes, solvents, co-solvents, and co-factors for use with existing force fields for proteins and other biopolymers. This can be done by splitting the target system into components, applying a more established force field (such as a protein force field) to any polymers present(such as a protein) and the selected SMIRNOFF force field to the other components of the system, then merging the results. This can easily be done via ParmEd; an example is provided in the `openforcefield` GitHub repository at https://github.com/openforcefield/openforcefield/tree/master/examples/mixedFF_structure, as well as in our SI.

### 5.4 SMIRNOFF provides a starting point for exploring development of next-generation force fields

SMIRNOFF force fields are only in their infancy, as we have validated the format and produced a simple general-purpose small molecule force field (SMIRNOFF99Frosst). While these are significant achievements, much more work is needed. We hope the community will put existing general purpose force fields in this format, resulting in improved extensibility (because chemical perception will no longer be hard-wired). Additionally, if our experience with AlkEthOH is any guide, this will likely result in uncovering and fixing many bugs.

We plan to build on our SMIRNOFF99Frosst prototype by refitting the parameters, first in a relatively conventional manner, which will likely improve accuracy. But refitting via novel methods will, in our view, be particularly exciting. We plan to utilize Bayesian techniques to refit SMIRNOFF force fields (e.g. as proposed in [119-121]) while varying both chemical perception and sampling over parameter space, which will open up exciting new lines of inquiry.

Overall, our hope is that SMIRNOFF will enable significant new force field science by reducing the amount of human expertise required as an *input* to force field parameterization. Direct chemical perception is enough simpler that that modification of SMIRNOFF force fields by non-experts becomes much more feasible than extension of force fields using traditional atom-typing.

## 6 Discussion and conclusions

In our view, the atom typing typically employed by fixed charge force fields introduces a variety of complexities and deficiencies in the force field development process. It tends to result in an explosion of parameters, a large potential for error (human or otherwise), and convolutes parameters making extension extremely difficult. At the same time, it also obfuscates the underlying chemistry and requires considerable human expertise to even understand the atom typing, much less manipulate it. It also makes automated fitting of force fields (one long-term goal of our effort) difficult, because a key input — the atom typing employed — is decided in advance by human experts.

As we have argued here, direct chemical perception simplifies the process of parameter assignment, reduces the number of force field parameters required, and helps reduce the potential for severe error both because of a more direct use of chemical information and because it allows for easy use of generic parameters. Thus as we showed here, shifting to direct chemical perception such as that implemented in the SMIRNOFF format immediately fixes several problems encountered by even modern fixed charge force fields, even without any specific attention paid to these problems and without any refitting of parameters.

We have provided a prototype force field, SMIRNOFF99Frosst, which is a logical descendant of the AMBER family force field parm@frosst [87] but covers a vast chemical space with less than 350 lines of parameters as opposed to the thousands of lines of parameters used in GAFF and other modern force fields. While we expect that SMIRNOFF99Frosst is probably not as accurate in general as these force fields yet, our tests on densities and dielectric constants indicate it is at least not substantially worse, and extensive validation on FreeSolv hydration free energies indicates roughly comparable performance. Thus, we believe this force field will provide a good starting point for future development, especially in view of its relative simplicity. It also takes advantage of a number of the key features provided by the SMIRNOFF format (version 0.1 of which is described here).

The SMIRNOFF format opens up a variety of avenues for further inquiry. We briefly highlighted how the use of partial bond orders can reduce complexity for bond stretching terms, but it will likely be worthwhile to see whether the same approach will carry over to other terms in the force field. For example, when dealing with nitrogen pyramidalization as discussed above, perhaps the partial bond order can serve to allow intermediate degrees of pyramidalization between those recognized by the force field via improper terms which depend on the partial bond order. Thus, rather than encoding SMIRKS patterns to account for how the presence of a distant electron-donating or electron withdrawing group modulates the geometry at a nitrogen center, simple local properties like partial bond order could be used to interpolate parameters with more minimal SMIRKS patterns. Possibly a similar approach can be used to reduce the complexity of torsions. Additionally, the use of direct chemical perception means that chemical perception could be modified as part of a parameterization effort, allowing greater automation of parameterization. We will explore these issues in future work.

Further work is needed to improve, extend, and develop the SMIRNOFF99Frosst force field, to bring other force fields into the SMIRNOFF format, and to make new force fields which more fully take advantage of the format and, perhaps, utilize automatic parameterization. Community participation in this effort — by putting existing force fields into this format, by testing SMIRNOFF99Frosst, or by helping to advance new avenues of inquiry enabled by the format — is welcomed.

## 8 Acknowledgements

DLM and CCB appreciate the financial support from the National Science Foundation (CHE 1352608) and the National Institutes of Health (1R01GM108889-01) and computing support from the UCI GreenPlanet cluster, supported in part by NSF Grant CHE-0840513. JDC appreciates support from the Sloan Kettering Institute, the National Institutes of Health (National Cancer Institute Cancer Center grant P30 758 CA008748), and the National Science Foundation (CHE 1738979). MKG appreciates support from NIGMS grant R01GM061300. MRS appreciates the support of the National Science Foundation (CHE-1738975).

We appreciate helpful discussions with Christopher Fennell (Oklahoma State), Bryce Manubay (University of Colorado), Paul Nerenberg (California State University, Los Angeles), and Patrick Grinaway (MSKCC). The authors are especially grateful to members of the computational chemistry community producing high-quality open source software, especially Jason Swails (ParmEd), without which this project would not be possible. We also particularly appreciate OpenEye Scientific Software for granting us an academic license, and for helpful discussions and support (including CIB’s sabbatical in the Mobley lab at UCI to help initiate this work).

## Footnotes

1 OpenEye’s OEChem Toolkit implementation is a modified version of the original MDL model. Originally, the MDL model considered two types of aromatic groups: six-membered rings with alternating single/double bonds including the perimeter bonds in azulene, and five-membered rings with two double bonds and hetero atoms or a carbon anion at the ring apex. The OpenEye implementation uses a more limited definition of aromaticity including only the former group. That is to say, only six-membered rings with alternating single and double bonds and azulene are considered aromatic. The five-membered rings then have formally assigned single or double bonds.

